# Unified function of FACT in mammalian chromatin replication and transcription, dissolving and restoring nucleosomes to counteract genome aggregation

**DOI:** 10.64898/2026.02.25.707725

**Authors:** R. Ciaran MacKenzie Frater, Rikke Rejnholdt Jensen, Joaquim Ollé López, Valentin Flury, Jeroen van den Berg, Anton Bournonville, Nicolas Alcaraz, Simai Wang, Hannah Richter, Marty Yang, Qian Du, Alexander van Oudenaarden, Vijay Ramani, Nils Krietenstein, Anja Groth

## Abstract

Nucleosomes with their associated modifications underlie genome organisation and regulation. Replication and transcription require nucleosome disassembly to access the DNA template. How this is orchestrated without jeopardizing chromatin function remains unknown. Here, we reveal a general, global requirement of the histone chaperone FACT in mammalian replication, transcription and chromatin maintenance. Upon acute FACT depletion, replisome and RNA polymerase progression is halted genome-wide and chromatin structure in their wake collapses with reduced nucleosome occupancy, irregular spacing and intermediate assemblies. Chromatin states deteriorate as modified histones are lost due to a lack of histone recycling. Chromatin fiber disorder further manifests in the 3D genome, triggering active genes to coalesce in aberrant microcompartments. This establishes a unifying mechanistic basis for mammalian chromatin replication and transcription, with FACT mediating nucleosome disruption and re-assembly and thereby guarding against spurious chromatin aggregation. Nucleosome organization can therefore dynamically regulate genome architecture.

## Main Text

Nucleosomes are the basal layer of genome organization and packaging, and their composition, organization, and modifications define chromatin states that regulate transcription and specify cell identity^1^. Nucleosomes also present a barrier^2^ that must be displaced in front of replisomes and RNA polymerase to gain access to the underlying DNA template^3–5^. The ability to maintain and propagate chromatin states in the face of these disruptive events is a fundamental property of eukaryotic life broadly relevant to human development, disease, and aging.

Histone chaperones can mediate both nucleosome disassembly and re-deposition of the released histones (recycling) and de novo nucleosome assembly from new histones to maintain and restore the chromatin landscape. They constitute a diverse group of proteins that bind histones and regulate histone interactions with DNA and other molecules^6,7^. Many histone chaperones have been implicated in histone dynamics during transcription^8–13^ and replication^14–23^, and in both settings, recycling defects might be compensated by de novo histone deposition^24–26^. It therefore remains a challenge to study nucleosome barrier effects and the importance of histone recycling in transcription and replication, especially in mammalian cells where tools for rapid genetic manipulation have been limited.

The FACT complex^27–29^, consisting of the SPT16 and SSRP1 subunits in mammalian cells^30^, is a particularly versatile histone chaperone capable of engaging all core histones, subnucleosomal structures, and whole nucleosomes in a variety of conformations^31–38^. FACT is proposed to aid in the disruption of nucleosomes and mediate histone recycling during both transcription and replication^29^. FACT was named for its ability to enable RNA Pol II transcription through chromatin in vitro^39–41^, and recently, yeast FACT was also shown to facilitate replication of chromatin templates in vitro^42,43^. Structural studies and modelling have provided a mechanistic understanding of FACT function in both processes by visualizing FACT engaged in nucleosome disruption and histone H3-H4 and H2A-H2B transfer during RNA Pol II progression and in front of the yeast replisome^44–47^. In yeast, the Pob3 and Spt16 subunits of FACT are essential and similarly, mammalian SSRP1 and SPT16 are common essential genes (dep.map^48^). Yeast genetics has established that FACT is required to maintain nucleosomes during transcription^8,24,49–55^, which protects against spurious transcription^8^. Genetic evidence also implicates yeast FACT in replication-coupled recycling of parental histones to the lagging strand^44,56^. However, the roles of FACT in chromatin maintenance during mammalian replication and transcription remain debated. Depletion and knockout studies across different cell types have reported varying effects of FACT loss on replication and cell proliferation^57–60^, and many have observed gene expression changes and reduced nucleosome occupancy in transcribed regions^58,60–62^. Yet, a consensus regarding the role of FACT is missing, possibly due to chaperone redundancies and adaptation. However, recent work, exploiting rapid and targeted degradation of SSRP1 in human cancer cells, demonstrated a global requirement for FACT in transcriptional elongation and RNAPII pausing by maintaining the +1 nucleosome^62^.

Here, we leverage the power of rapid degron technology to address the acute consequences of SPT16 loss in mouse embryonic stem cells (mESCs) using a suite of advanced technologies to measure RNA and DNA synthesis genome-wide, and chromatin organization from nucleosome occupancy and modifications to genome architecture. We find that both replication and transcription arrest abruptly and globally as replisome and RNA Pol II progression is halted. In the wake of both machineries, chromatin is severely disordered with low nucleosome occupancy, irregular fiber organization and intermediate nucleosome assemblies. This argues for a unifying role of FACT in mammalian transcription and replication, removing nucleosome barriers and recycling histones to maintain nucleosome organization. In transcribed regions, the lack of FACT resulted in a collapse of the active chromatin state, with a drop in promoter and gene body modifications exceeding the overall nucleosome loss. Therefore, retention of pre-existing histones through transcription is critical to uphold high levels of active histone modifications – de novo incorporated H3.3 failed to compensate. These defects in the primary chromatin structure impacted genome contacts and architecture in FACT-depleted cells. The interaction frequency between a large fraction of active genes increased dramatically, forming what we suggest to be a novel, aberrant type of micro-compartments. Across a suite of genomic, chromatin and functional features, the strong loss of ordered nucleosomes in TSS proximal regions was predictive for aberrant microcompartment (AMC) formation. This argues that well-ordered nucleosome fibers – maintained through recycling of pre-existing histones – counteract chromatin aggregation.

## Results

### FACT is an essential co-factor for DNA and RNA synthesis

To study the function of mammalian FACT, we used the dTAG system^63^ to generate a mESC model for the acute depletion of SPT16. We focused on SPT16, as it is the primary subunit responsible for H3-H4 engagement in recent structures of chromatin replication and transcription intermediates^44,46^. This resulted in a strong depletion of SPT16 within 1 hour of dTAG treatment (Figures 1A and S1A). SPT16-depleted cells showed reduced viability by 6 hours and most of the cells died within 24 hours (Figure 1B), indicating that mESCs do not tolerate loss of FACT. To measure the rates of global replication and transcription, we used EdU and EU labelling followed by high-content microscopy, focusing on early timepoints after SPT16 depletion before viability was challenged. Global RNA and DNA synthesis declined rapidly and on the same timescale (Figures 1C, 1D, 1G, 1H, 1I, S1B, S1C, and S1D), reaching a minimum within 1 and 2 hours of SPT16 depletion, respectively. This was not associated with a reduction in S-phase cells, as the cell cycle distribution was largely unaltered up to 4 hours of dTAG treatment (Figures S1D and S1E). The stark arrest of DNA and RNA synthesis was not accompanied by DNA damage or activation of the replication checkpoint (Figures 1E and 1F). This demonstrates that mammalian FACT is a vital cofactor for transcription and DNA replication in mESCs.

**Figure 1.**
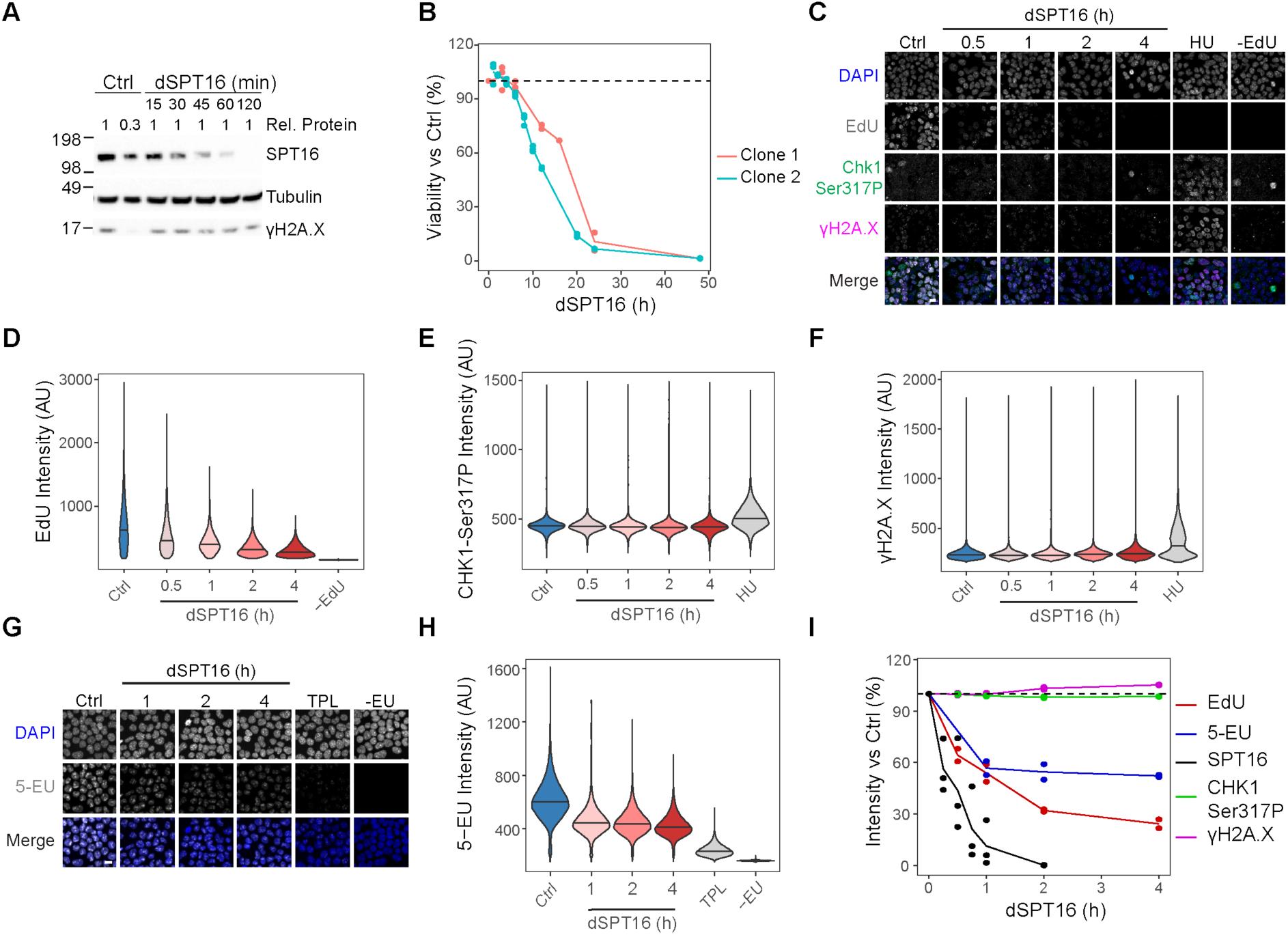
FACT is essential for global levels of replication and transcription. **A.** Western blot of SPT16, α-Tubulin, and γH2A.X in mESCs depleted for SPT16 by dTAG treatment (dSPT16) or treated with DMSO vehicle control (Ctrl) for the times indicated. The relative amount of extract loaded is indicated (Rel. protein). **B.** Cell viability of two SPT16-dTAG mESCs clones. Each point represents a biological replicate (n=2-3 per timepoint per clone). **C.** Representative images of EdU, γH2A.X and CHK1-Ser317P. Ctrl, hydroxyurea (2 mM for 4 hours, HU) treatment, and unlabelled (-EdU) cells were included as controls. Scale bar, 150 μm. Quantifications are shown in Figure 1D-F. **D.** Quantification of EdU intensity in EdU-positive from C. 27116 nuclei were measured per condition. Values are representative of 2 biological replicates. **E.** Quantification of mean CHK1-Ser317P signal from C. **F.** Quantification of mean γH2A.X signal from C. **G.** Representative images of 5-ethynyl uridine (5-EU) incorporation in SPT16-dTAG mESCs. Triptolide treatment (10 μM for 4 hours, TPL) and unlabelled (-EU) cells were included as a control. Scale bar, 150 μm. **H.** Quantification of mean 5-EU intensity from G. Values are representative of 2 biological replicates. 54608 cells were measured per condition. **I.** Graph indicating time dependent changes in SPT16 levels (western blot, A) and EdU (D), γH2A.X (E), CHK1-Ser317P (F), and 5-EU (H) relative to Ctrl. Each dot represents a distinct biological replicate (n=3 western blot, n=2 microscopy).

### FACT is required for progression of RNA and DNA polymerases in chromatin

To understand how replication is arrested and whether there is a connection to the arrest of transcription, we first assessed genome-wide replication patterns by EdU-seq after 1 hour of SPT16 depletion when replication was reduced but still detectable (Figure 1I). The EdU signal was reduced genome-wide across replication timing, showing a global requirement for FACT in DNA synthesis across S phase (Figure S2A), though early replication was less impaired. Transcribed genes, non-transcribed genes and intergenic regions showed a largely similar reduction in DNA synthesis (Figure S2B), arguing that the replication arrest is independent of problems with transcription. However, the genomic distribution of the EdU signal, representing replication activity, was altered upon FACT depletion (Figure S2C). EdU incorporation was only moderately reduced in initiation zones (IZs) but strongly declined outside of IZs (Figures 2A and 2B). This pattern points to impaired replication fork progression being responsible for the loss of DNA synthesis rather than an initiation defect. The less dramatic reduction in EdU incorporation in early-replicating regions (Figure S2A) likely reflects the higher number of IZs^64^. Indeed, comparing replication activity downstream of IZs to replication within IZs showed that replication elongation is globally impaired regardless of replication timing (Figure 2C).

**Figure 2.**
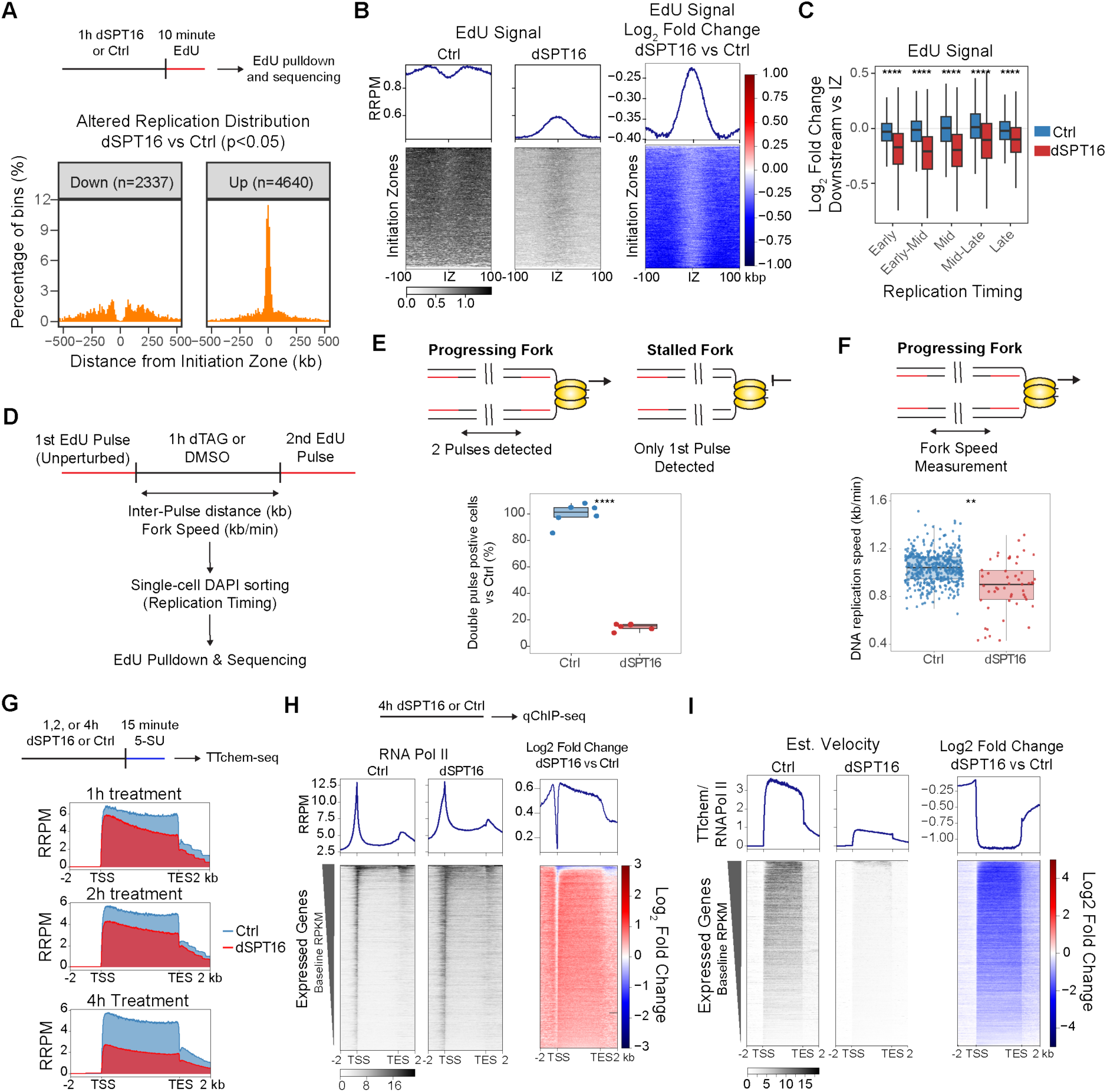
FACT promotes replisome and RNA PolII progression in chromatin genome-wide. **A.** Distribution of replication activity across the genome. Distance to the nearest initiation zone (IZ) for regions with significant changes in EdU distribution (RPM) in SPT16-dTAG mESCs after 1 hour dTAG (dSPT16) relative to DMSO vehicle control (Ctrl). **B.** Heatmap of EdU signal (as in A) quantified by spike-in over initiation zones (IZs). Spike-in normalisation was performed as reference-adjusted reads per million (RRPM). **C.** EdU signal quantified by spike-in (as b) in 75 kb bins centred at IZs or 150 kb downstream. Individual IZs were stratified by replication timing. **D**. Schematic of scEdU-seq to capture ongoing replication in SPT16-depleted mESCs. **E.** Proportion of cells with measurable double EdU pulse signal normalized to DMSO control. Each dot represents the mean of a technical replicate of cells in a 384-well plate (n=6). **F.** DNA replication speeds in kilobases per minute (kb/min) in cells with detectable double pulse signal. **G**. Median TTchem-seq signal of expressed genes normalized by spike-in. **H**. Spike-in normalised RNA Pol II occupancy across expressed genes shown as RRPM. Heatmaps are sorted by baseline gene expression rate with profiles of mean signal above. **I**. Estimated velocity of all expressed protein-coding genes after 4 hours of dSPT16 or Ctrl treatment. Estimated velocity represents the ratio of nascent RNA production (TTchem-seq signal) to the amount of RNA Pol II present (qChIP-seq signal).

To directly address the requirement for FACT in replication elongation, we analyzed replication fork dynamics in single cells by scEdU-seq where fork progression is measured by double EdU pulse labelling. dTAG was added after the first EdU pulse to compare progression in the presence and absence of SPT16. This setup also enables quantification of other DNA replication metrics^65^ such as fork number and replication timing (Figure 2D). We captured similar numbers of replication forks per cell independent of replication timing, and DNA replication timing was not altered after FACT depletion (Figures S2E, S2F, and S2G). Therefore, FACT depletion does not lead to substantial new origin firing that perturbs replication timing patterns. However, over 80% of the cells lacked double pulse signal upon SPT16 depletion (Figures 2E and S2H), implying that FACT is essential for replication fork progression in mESCs. In the residual 20% of SPT16-depleted cells where fork progression was still detectable, fork speeds were significantly slower (Figure 2F) and fork slowdown was comparable across S phase (Figure S2I). This demonstrates that ongoing replication forks, irrespective of genomic context, were unable to progress through chromatin in the absence of FACT, supporting a role of FACT as a general replisome progression factor. We conclude that without FACT, replication can initiate but replication forks arrest in a stable checkpoint-blind state genome-wide, suggestive of a block to CMG (Cdc45-MCM-Gins) unwinding activity by nucleosomes.

To address the requirement for FACT in transcription in comparison to replication, we performed TTchem-seq^66^. Global RNA Pol I and II transcription was progressively reduced upon SPT16 loss (Figures S2J, S2K, S2L and S2M), as reported for SSRP1 depletion in leukaemic cells^62^. More than 3000 and 10,000 genes were downregulated after SPT16 depletion for 1 and 4 hours, respectively, which is substantially more than reported for SSRP1 depletion (657 and 3,703 genes at 1 and 4 hours, respectively^62^). Highly transcribed and longer genes were most affected by SPT16 loss (Figures S2N and S2O), likely explaining why transcription of the very short RNA Pol III genes does not require FACT. Furthermore, metagene analysis showed a 5’ bias in transcription levels (Figure 2G), most prominent at longer genes (Figures S2P, S2Q, and S2R), consistent with defective transcription elongation as reported for SSRP1 depletion^62^. ChIP-seq analysis calibrated by spike-in showed strong RNA Pol II accumulation across gene bodies and even upstream of the TSS (Figure 2H). The combined reduction in nascent RNA synthesis and increase in elongating polymerase led to a strong reduction in estimated transcription velocity (Figure 2I). Therefore, there is an accumulation of slow-elongating RNA polymerases in the absence of FACT, as progression through chromatin is hindered by nucleosomes ahead of the RNA polymerase complex. In combination with the replication fork progression defect, this highlights the requirement for FACT as a general chromatin disassembly factor, facilitating the efficient progression of macromolecular replication and transcription machines through chromatin.

### FACT maintains nucleosome structure and organization through replication and transcription

To measure the contribution of FACT-mediated histone recycling to chromatin restoration during replication and transcription, we performed replication-aware single-molecule accessibility mapping (RASAM)^67,68^ that globally and non-destructively measures protein-DNA interactions on individual nascent and bulk chromatin fibers. The abundance of BrdU positive (BrdU+) fibres was reduced in FACT-depleted samples as expected from our microscopy data (Figure S3A) and BrdU+ fibres originated from regions closer to initiation zones (Figure S3B and S3C), similar to EdU-seq (Figures 2A, 2B and 2C). Nucleosome footprints in BrdU+ fibres also showed the expected shift towards smaller footprint sizes compared to BrdU-fibres (Figure S3D)^67^. Therefore, RASAM captures chromatin fibres replicated under SPT16-depleted conditions. Nascent dSPT16 fibers showed reduced nucleosome occupancy (Figures 3A and S3E) and more fibers had irregular nucleosome spacing, which correlated with increased accessibility (Figures 3B, 3C, S3F, S3G, and S3H). This pattern resembles nascent fibers replicated in the absence of CAF-1^67^, which is required for the de novo deposition of new histones. In the absence of SPT16, there was also an increase in smaller footprints on nascent chromatin, with sizes consistent with hexasomes, tetrasomes, and dimers (Figures 3D and S3I). To understand which nucleosomal structures were enriched, we applied Iteratively Defined Lengths of Inaccessibility (IDLI)^69^ (Figures S4A and S4B). Nascent chromatin fibers from dSPT16 cells (Figure S4C) were enriched for four types of nucleosomes, all with smaller footprints and higher accessibility (Figures 3E and 3F). This confirmed enrichment of subnucleosomal structures such as hexasomes, tetrasomes, and dimers, while fully assembled nucleosomes were depleted (Figure S4D). The two most highly enriched clusters showed accessibility patterns consistent with tetrasome and hexasome footprints (Figures S4C and S4D). Together, this suggests that fewer intact nucleosomes are assembled on nascent chromatin and that if H3-H4 tetramers are assembled, there is an additional defect in H2A-H2B deposition. Together, RASAM demonstrates that mammalian FACT is essential for maintaining the occupancy of all core histones through replication.

**Figure 3.**
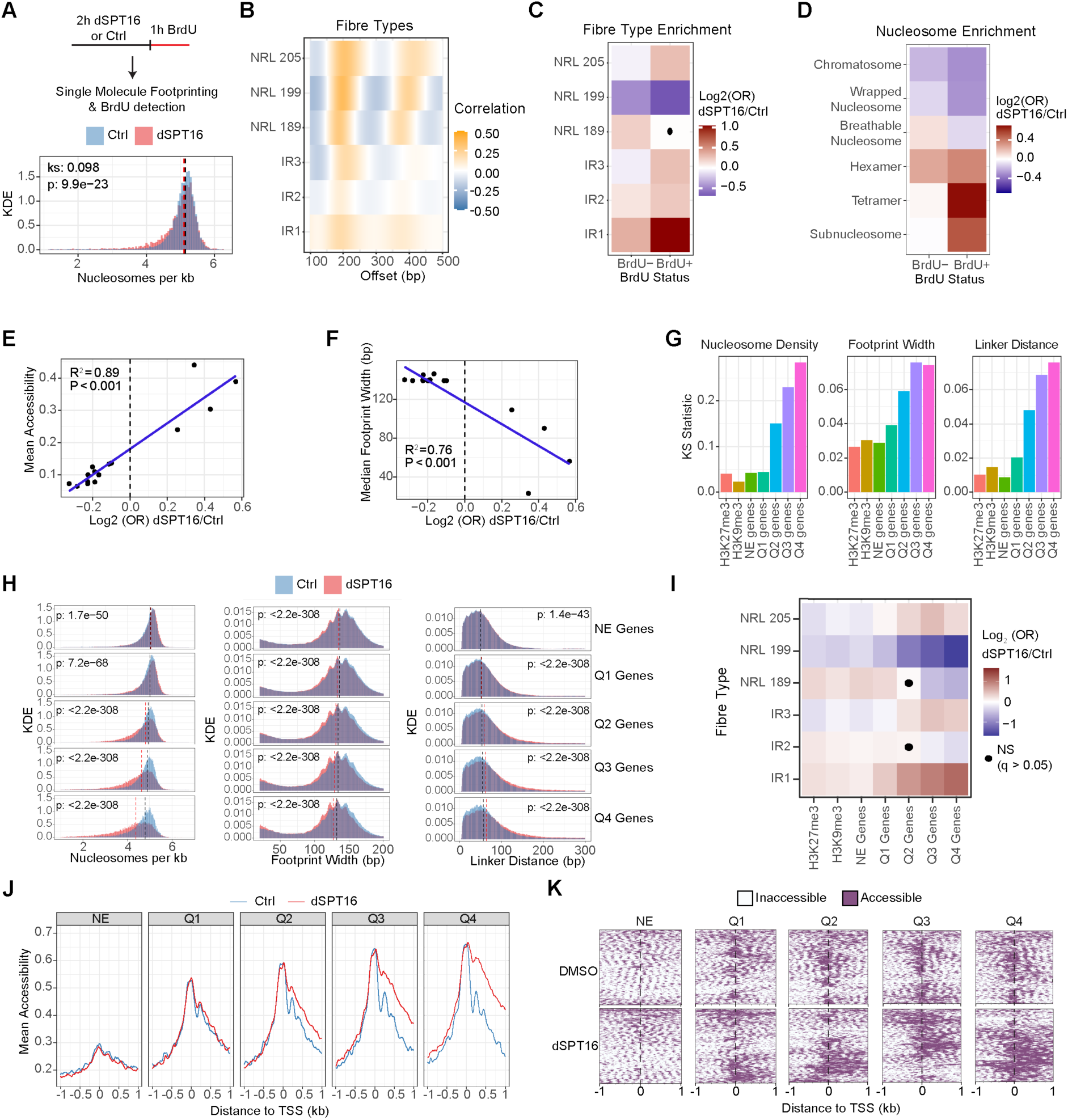
Histone recycling maintains chromatin fibre structure during replication and transcription. **A.** (top) Experimental setup for RASAM, BrdU was added to SPT16-dTAG mESCs after 2 hours of treatment with dTAG (dSPT16) or DMSO vehicle (Ctrl). (bottom) Distribution of single-molecule nucleosome density estimates on nascent chromatin fibres. Dotted lines represent the median values for each condition, Kolmogorov-Smirnov effect size (ks) and p value are shown. **B.** Visualisation of average fibre types detected by autocorrelation (NRL: nucleosome repeat length, IR: irregular). **C.** Heatmap of log_2_ odds ratios (OR) showing enrichment (red) or depletion (blue) of fibre types on bulk and nascent chromatin after dSPT16 treatment. Significance was determined by Fisher’s exact test, and non-significant (p < 0.05) conditions are marked by a dot. **D.** Relative enrichment (red) or depletion (blue) of nucleosome structures in bulk and nascent chromatin upon dSPT16. Nucleosome structures are based on footprint widths shown in Figure S3I. All conditions were significantly changed based on Fisher’s exact test. **E,F.** Mean accessibility (**E**) and footprint width (**F**) of nucleosome types plotted against their enrichment in dSPT16 nascent fibers. Nucleosome types were detected by IDLI (Figure S4D) and enrichment calculated as Log_2_ OR of dSPT16 versus DMSO. Pearson correlation is shown. **G** Kolmogorov-Smirnov (KS) effect sizes showing the magnitude of the difference in nucleosome density, footprint widths, and DNA linker distances between dSPT16 and Ctrl BrdU-fibres overlapping H3K9me3 or H3K27me3 regions, non-expressed (NE) genes, or genes ranked by quartile of expression (increasing from Q1 to Q4). **H.** Distributions of nucleosome density, footprint width, and DNA linker distance in dSPT16 and control fibres Dotted lines represent the median values of Ctrl (black) and dSPT16 (red) fibres. P-values from KS test are shown. **I.** Heatmap of fibre type enrichment as in **C** for fibres overlapping the specified annotations. **J.** Mean modification probability of all fibres covering TSS of genes stratified by expression levels (as in **G,H**). **K.** Single-molecule accessibility patterns of n=500 randomly-sampled chromatin fibres from J. The dotted line represents the TSS and each line on the y-axis represents the modification status of a single fibre overlapping the region.

To study the effects of SPT16 depletion on transcription-coupled chromatin maintenance, we analysed the BrdU negative (BrdU-) fibres covering genic regions from RASAM. Similarly to replication, transcription in the absence of SPT16 caused reduced nucleosome occupancy and smaller footprint sizes, with changes scaling with transcription rate (Figures 3G, 3H and S4E). Heterochromatin regions did not show reduced nucleosome occupancy but had a slight reduction in footprint width (Figure S4F), consistent with a role for FACT in maintaining H2A/H2B occupancy in heterochromatin^70^. Nucleosome organization was also impaired by transcription in the absence of FACT, with an increase of longer inter-nucleosome DNA linkers and irregularly spaced chromatin fibres (Figures 3G, 3I, S4G, and S4H). Chromatin disruption in transcribed genes began immediately downstream of the TSS and extended substantially into the gene body (Figure 3J), with the appearance of long nucleosome-depleted regions evident on single chromatin fibres (Figure 3K). We conclude that FACT is a universal histone recycling factor for replication and transcription, restoring nucleosome fiber structure in the wake of polymerases.

### Histone recycling maintains chromatin composition and markup

To reveal potential broader implications of histone recycling for chromatin integrity, we performed spike-in corrected ChIP-seq (qChIP-seq) in FACT depleted cells focusing on transcription-associated modifications and histone variants. Of note, the strong arrest of DNA replication precluded a broader analysis of post-replication chromatin restoration. Transcription in the absence of FACT led to a strong reduction in the occupancy of core histones H2B and H3 at the TSS and TSS-proximal regions of active genes (Figures 4A and 4B), indicative of substantial nucleosome loss consistent with our single-molecule data (Figures 3J and 3K) and previous observations in yeast^24,50,53^. Concomitantly, H3.3 occupancy increased at TSSs and throughout gene bodies in a manner scaling with transcription rate (Figures 4A and 4B), arguing that H3.3 deposition is increased to compensate for nucleosome loss when FACT-mediated recycling is disrupted. The histone recycling defect was accompanied by a marked loss of gene body modifications associated with active transcription, H3K36me3 and H2BK120ub1, in a manner scaling with transcription rate (Figures 4C and 4D). Furthermore, the active promoter modifications H3K4me3 and H3K27ac were strongly depleted, again with stronger changes at the most highly transcribed genes (Figures 4E and 4F). H2A.Z, which is present in both active and inactive genes in euchromatin^71^, was also depleted around the +1 nucleosome of highly transcribed genes (Figures 4E and 4F), but the occupancy increased downstream of the TSS (Figure 4F). This is reminiscent of reports of altered Swr1 activity upon loss of FACT in yeast^55^. The irregular spacing of nucleosomes (Figures 3H and 3I) might cause Swr1 (in mammals SRCAP) mislocalization, as it measures nucleosome linker length to engage with the +1 nucleosome^72^. Notably, the loss of histone modifications was substantially more pronounced than the loss of total histones (Figure 4G). This argues that although compensatory H3.3 deposition partially restores nucleosome occupancy across the gene, it cannot substitute for transcription-coupled recycling in maintaining high levels of active histone modifications. Collectively, these results demonstrate that FACT-mediated transcription-coupled histone recycling is vital to maintain the chromatin state of active genes.

**Figure 4.**
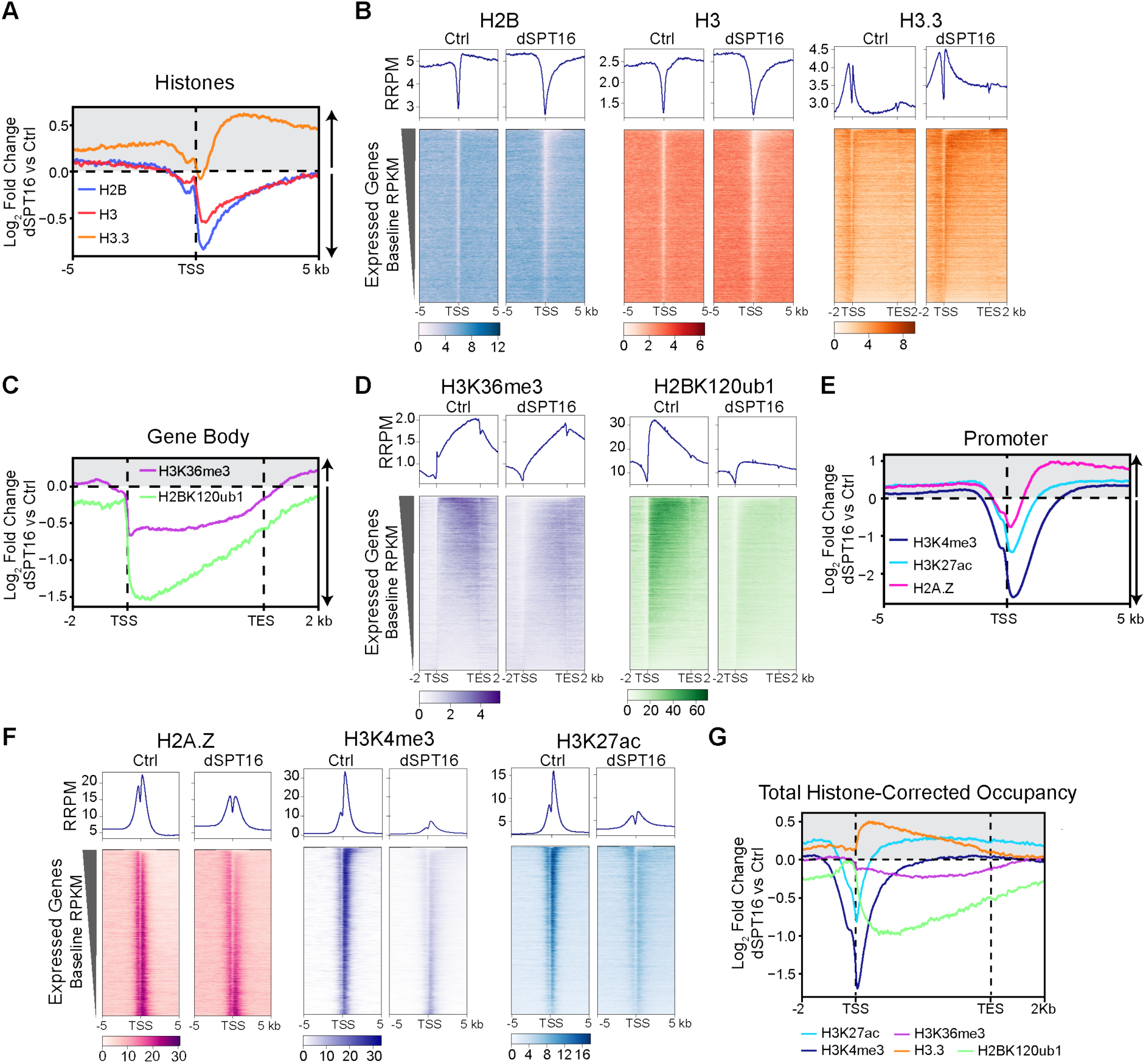
Histone recycling is required to maintain the active chromatin landscape. **A.** Log_2_ fold change of histone H2B, H3, and H3.3 occupancy at transcription start sites (TSS) of highly transcribed genes (top 25% of expression, Q4) comparing dSPT16 (dTAG, 4 hours) to DMSO (Ctrl) treated cells. Arrows indicate increased and decreased occupancy upon FACT depletion. **B.** Heatmaps of histone H2B, H3 and H3.3 occupancy shown as RRPM normalised signal across all expressed protein-coding genes sorted by descending baseline transcription rate. Profiles represent the mean signal at each position. **C.** Log_2_ fold change of H2BK120ub1 and H3K36me3 levels across gene bodies shown as in A. **D.** Heatmaps of H2BK120ub1 and H3K36me3 levels across expressed protein-coding genes as in B. **E.** Log_2_ fold change of H3K4me3, H3K27ac, and H2A.Z levels at TSSs as in A. **F.** Heatmaps of H3K4me3, H3K27ac, and H2A.Z levels as in B. **G.** Log_2_ fold change occupancy of active modifications across Q4 genes after normalisation to H2B (H2BK120Ub) and H3 (H3K27ac, H3K4me3, H3K36me3, and H3.3) signal. (A-G) All ChIP signals were spike-in normalized. RRPM, reference-adjusted reads per million.

### Active genes coalesce into aberrant micro-compartments upon loss of FACT

To assess how chromatin maintenance by FACT shapes genome organization, we performed Micro-C^73,74^ over a time course of 1, 2, and 4 hours of SPT16 depletion. Interaction maps revealed a striking appearance of novel interaction features between genes that resemble micro-compartments in form and shape (Figure 5A)^75–77^. Visual inspection suggested that these aberrant micro-compartments (AMCs) form between genes and increase in strength over time with SPT16 depletion (Figures 5A and S5A). Comparing interaction heatmaps to eigenvalues at 5 kbp resolution showed increased eigenvalues in the regions with higher interaction frequency (Fig. 5A). Focal interactions, that appear as dots in heatmaps, emerge predominantly from convergent CTCF-CTCF and from enhancer-promoter (E-P) and promoter-promoter (P-P) interaction sites^74,78–82^. Micro-compartments are reported to form primarily between cis-regulatory elements, including enhancers^75–77^. To investigate the relationship between AMCs and known elements that form focal interactions or micro-compartments genome-wide, we performed pile-up analysis at putative CTCF-CTCF, E-P, and P-P interaction sites ranging from 10 to 500 kb and stratified them into equally sized groups (Figure 5B). Upon SPT16 depletion, P-P interactions increase dramatically, while E-P interactions remain unaltered and CTCF-CTCF interactions decrease moderately. The latter appeared unrelated to perturbation of transcription in SPT16-depleted cells, as stratification of convergent CTCF-CTCF interactions by TSSs proximity did not reveal differences (Figure 5C). However, Cohesin occupancy increased moderately at CTCF sites, suggesting a slightly enhanced barrier function in the absence of FACT (Figure S5B). Together, these data link AMCs to P-P interactions and argue that FACT is not directly required for cohesion-mediated loop extrusion, which sustains CTCF-CTCF and EP interactions^78,83–86^.

**Figure 5.**
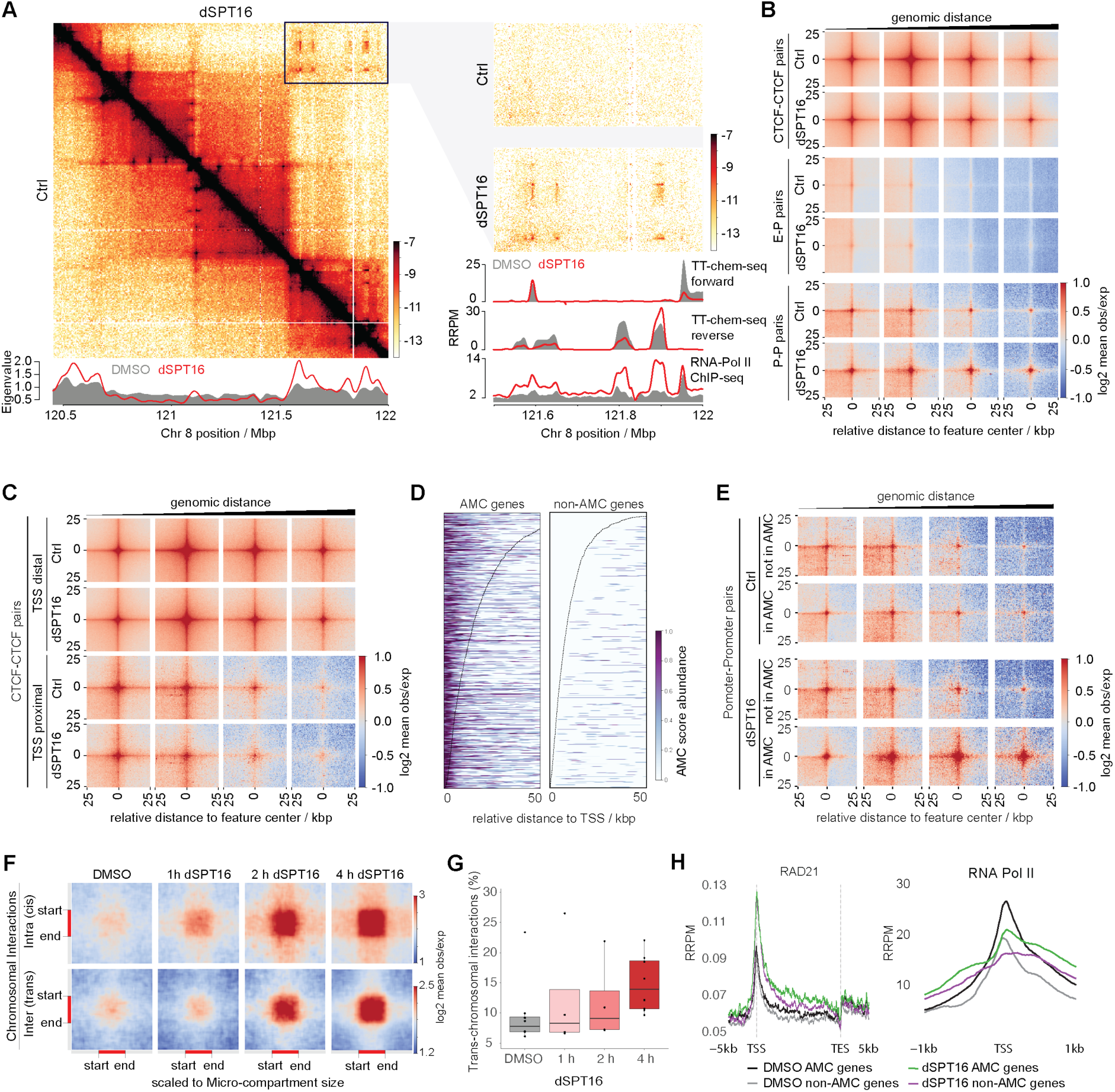
FACT counteracts affinity-driven coalescence of active genes into AMCs. **A.** Micro-C interaction heatmaps (left: 10 kbp, right: 2 kbp resolution) of cells treated for 4 hours with dTAG (dSPT16) or DMSO (Ctrl). AMC formation correlates with increased eigenvalues (signal tracks, bottom left). TTchem-seq and RNA Pol II qChIP-seq data tracks expressed in reference-adjusted reads per million (RRPM) (right, bottom). **B.** Pile-up analysis of all putative CTCF-CTCF, enhancer-promoter (E-P), and promoter-promoter (P-P) contacts at 500 bp resolution with a 25 kbp flank. Putative pairs were computed for 10 to 500 kbp distances and stratified into four equally sized bins (left to right). **C.** As B, but for CTCF-sites stratified by co-localization with TSS or not. **D.** Abundance of AMC called bins over AMC and non-AMC gene bodies aligned at TSS. **E.** same as B, but for P-P intersections stratified by associations with AMCs or not. **F.** Cis and trans chromosomal domain interaction pile-ups for regions assigned as AMCs. **G.** Fraction of trans chromosomal interactions. **H.** ChIP-seq profile for RAD21 and RNA Pol II at AMC-forming and non-forming genes. ChIP signals were spike-in normalized. RRPM, reference-adjusted reads per million.

To identify the genomic positions of AMCs, we developed an algorithm that defines AMCs based on eigenvalue increase after SPT16 depletion (see methods, Figure S5C). We detected a total of 5057 AMCs genome-wide, with a mean size of ∼15 kb. In agreement with our observation that AMCs correlate with genes, 4412 out of 19365 genes intersected with AMCs. Notably, within the top 25% of expressed genes, approximately half of the genes coalesced into AMCs in the absence of FACT (Figure S5D). The number of genes located in AMCs was reduced with lower expression levels and fewer non-expressed genes formed AMCs. Across individual genes, the TSS-proximal genic regions were most prone to form AMCs (Figure 5D). To confirm our computational approach to detect AMCs, we compared the interactions between putative pairs of TSSs of the top 25% expressed AMC and non-AMC genes in the range of 10 to 500 kb (Figure 5E). The pile-up analysis showed that P-P interactions specifically increased for AMC genes and not for non-AMC genes, validating our computational AMC classification.

### AMC formation involves affinity-driven self-association of active genes

Micro-compartmentalization is proposed to be an affinity-driven process as it is resistant to Cohesin perturbation^75^. During the mitosis-to-G1 transition, micro-compartments are formed by cis-regulatory elements, including enhancers and promoters marked by H3K27ac^76,77^. Perturbations of transcription, including loading, elongation, and degradation of RNA Pol II, have been shown to affect interactions between cis-regulatory elements^87,88^, suggesting that affinities between Polymerases may drive genes to coalesce^89^. To assess whether AMC formation is loop-extrusion- or affinity-driven, we performed cis- and trans-chromosomal compartment pile-up analysis. We observed that the AMC interactions, inter- and intra-chromosomal alike, increased dramatically with the time of SPT16 depletion (Figure 5F), as did global trans-chromosomal interactions (Figure 5G). This supports the idea that AMC formation is affinity-driven rather than a loop-extrusion-driven process, which operates in a cis manner.

Although AMCs association scaled with transcription rate, RNA Pol II activity did not appear to be the determining factor, as only half of the top 25% of transcribed genes formed AMCs. To evaluate roles of RNA Pol II binding and loop extrusion in AMC formation, we compared qChIP-seq profiles of RNA Pol II and RAD21 of the top 25% AMC and non-AMC-forming genes (Figure 5H). AMC-forming genes had slightly higher levels RNA Pol II and Cohesin in the control condition and after SPT16 depletion, but the differences were modest. The limited correlation of RNA Pol II and Cohesin accumulation with AMCs was also evident from visual inspection of qChIP-enrichment with AMC location (Figure 5A, right). Therefore, RNA Pol II accumulation and Cohesin alone do not explain the coalescence of a subfraction of active genes into AMCs.

### Irregular and disassembled nucleosome fibers underlie AMC formation

To identify features of FACT-depleted chromatin that could explain AMC formation, we developed a classification model to predict AMC formation based on chromatin alterations (qChIP after SPT16 depletion) GC content, exon density, proximity to IZs, gene density, and transcriptional activity (see methods). The model reliably predicted AMC-forming genes with about 75% accuracy (Figures S6A and S6B). Using SHAP analysis, we identified the features most predictive of AMC formation. Gene expression was most predictive for AMC formation (Figures 6A), consistent with these structures forming more often at highly transcribed genes (Figure S5D). However, less than 50% of highly expressed genes form AMCs (Figures S5D), underscoring that it must act in combination with other features in driving AMC formation. Other top features were lower nucleosome occupancy downstream of the TSS and lower H2A.Z levels after SPT16 depletion (Figures 6A and S6C). Unbiased clustering of SHAP values efficiently separated AMC genes from non-AMC genes (Figures S6D and S6E), validating their predictive value and again highlighting reduced nucleosome occupancy as the key predictive feature (Figures S6D and S6F). Other important features included lower gene body GC content, which might reduce nucleosome stability, lower gene density and higher H3K27ac and H2BK120ub1 levels after SPT16 depletion (Figures 6A and S6C).

**Figure 6.**
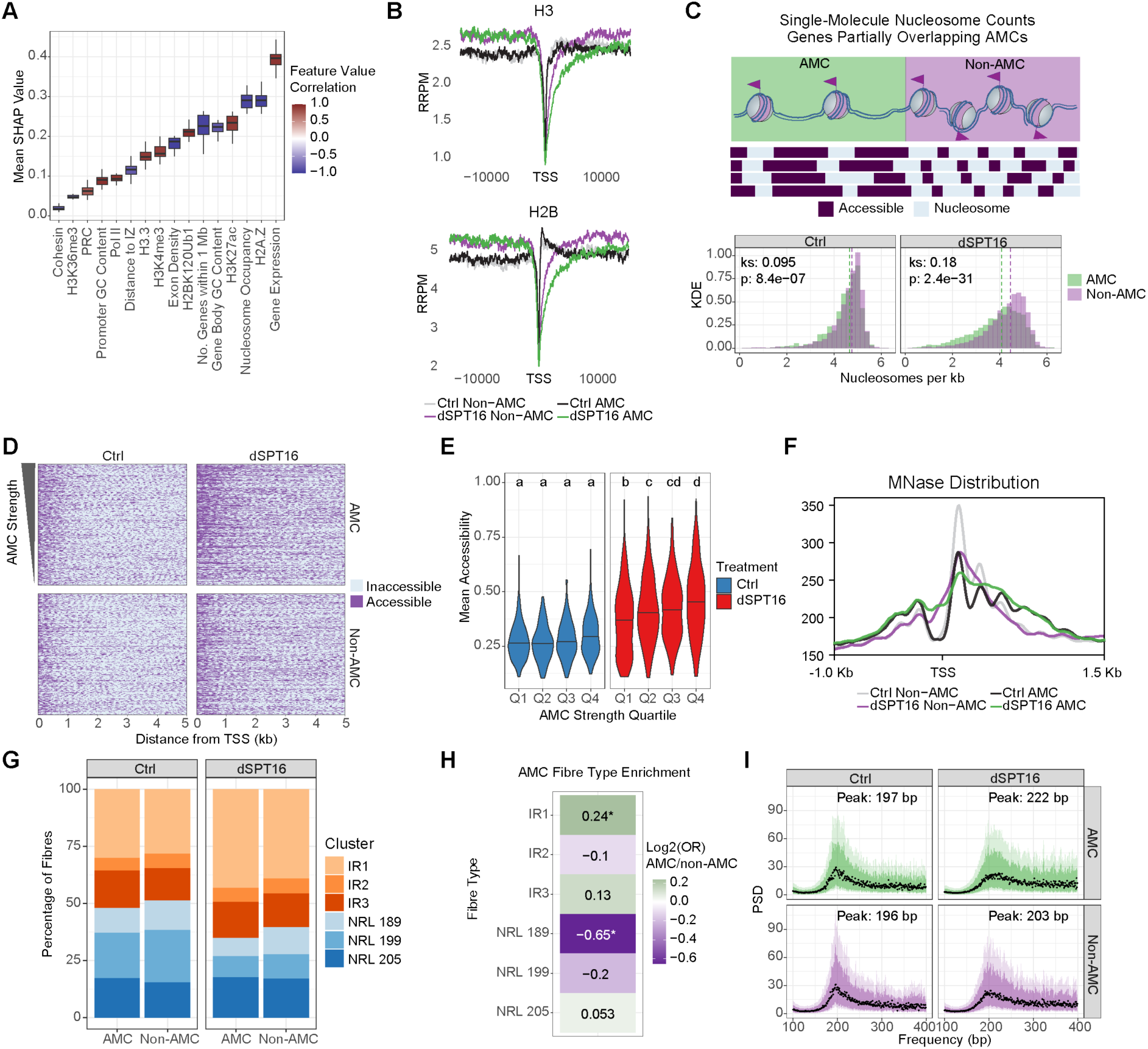
Collapse of chromatin fibre structure underlies AMC formation. **A.** Mean SHAP importances for each feature included in the model for predicting AMC formation. Boxplots represent mean SHAP values for n=16 cross-validation folds. Boxplot colour represents the correlation between feature values and a positive AMC prediction. **B.** Occupancy of H3 and H2B for expression-matched AMC and non-AMC genes (data from Figure 4A). RRPM, reference-adjusted reads per million. **C.** Nucleosome occupancy on single chromatin fibres overlapping AMC and non-AMC regions of individual genes (data from Figure 3G-K). **D.** Single chromatin fibres overlapping the first 5 kb of AMC or non-AMC genes. N=500 fibres were randomly sampled for each annotation and condition. AMC fibers are sorted by decreasing AMC strength. **E.** Mean accessibility across individual chromatin fibres covering the entire first 5 kb of AMC genes after DMSO Ctrl (n=792) or dSPT16 (n=834) treatment. Fibres were stratified into quartiles based on increasing AMC strength (Q4 highest). Significance was calculated by ANOVA followed by Tukey’s post-hoc test. Conditions containing the same letter were not significantly different (p > 0.05). **F.** MNase profile after Ctrl or dSPT16 treatment for 4 hours. **G.** Proportion of chromatin fibre types within the first 5kb of expression-matched AMC and non-AMC genes. n=2646 Ctrl AMC, n=4114 dSPT16 AMC, n=2443 Ctrl non-AMC, and n=3878 dSPT16 non-AMC. **H.** Relative enrichment of fibre types as in G. Odds ratio (OR) of enrichment was calculated by Fisher’s exact test (*: p < 0.05). **I.** Nucleosome repeat length variation. Power spectral density (PSD) of nucleosome repeat lengths measured by Fourier transformation of m6A patterns in RASAM reads overlapping the first 5 kb of expression-matched AMC or non-AMC genes (fibres from G). Black points represent the median PSD for each repeat length, darker shading represents the interquartile range (IQR), and lighter shading represents the IQR*1.5. The peak value shown represents the chromatin repeat length with the highest median spectral density.

To compare chromatin composition independent of expression levels, we generated an expression-matched set of AMC and non-AMC genes (Figure S6G). Among these, AMC genes showed higher levels of active modifications, including H3K27ac, H3K36me3 and H2BK120ub1 than non-AMC genes in unperturbed conditions (Figure S6H). Both AMC and non-AMC genes lost active modifications upon SPT16 depletion, with the former ending at a higher level of H3K27ac and H2BK120ub1 relative to non-AMC genes after SPT16 depletion. The predictive value of high H3K27ac and H2BK120ub1 levels thus reflects a higher modification state of these genes in unperturbed conditions rather than a specific increase in modification levels after FACT depletion. While it is unclear why this gene set has a higher modification state compared to their expression-matched non-AMC counterparts, our model indicates that this property contributes to their propensity for AMC formation. RNA Pol II was increased throughout the gene body of AMCs, arguing that this could contribute to AMC formation (Figure S6H), though it alone was a less important predictor than chromatin composition (Figure 6A). RAD21, SUZ12, and RING1B did not accumulate at AMCs, consistent with Cohesin and Polycomb complexes not driving AMC formation (Figure S6H). Importantly, AMC genes had greater loss of both H2B and total H3 in the early gene body where AMCs are enriched (Figure 6B), which supports that disrupted chromatin fibre structure could underlie AMC formation. H2A.Z was depleted downstream of the TSS, consistent with nucleosome loss. Concomitantly, H3.3 deposition was strongly increased across AMC genes – also in comparison to non-AMC genes – arguing that more severe defects in histone retention invoke elevated de novo deposition activity in an attempt to maintain nucleosome organization (Figure S6G).

To reveal how chromatin fibre structure is perturbed in AMCs, we compared single-molecule chromatin fibre accessibility for AMC and non-AMC regions using our RASAM data. Chromatin fibres overlapping both AMC and non-AMC regions of the same set of genes showed similar footprint widths (Figure S6H), but nucleosome occupancy was strongly reduced within the AMC regions (Figure 6C). To ensure that AMC detection is independent of changes in MNase accessibility in Micro-C experiments, we validated AMC formation by Hi-C analysis (Figure S6J). Chromatin fibres overlapping TSS proximal regions of expression-matched AMC and non-AMC genes showed a higher degree of disruption for AMC genes, with accessible DNA scaling with AMC strength (Figures 6D and 6E). MNase profiles further showed reduced nucleosome phasing (Figure 6F). This reflected increased disorder in nucleosome positioning, as the proportion of irregularly-spaced chromatin fibres was increased in AMCs and regularly-spaced fibres with shorter repeat lengths were depleted (Figures 6G and 6H). Furthermore, nucleosomal repeat length became more variable in AMCs after SPT16 depletion with predominantly longer spacing (Figure 6I). Collectively, these lines of evidence suggest that the loss of nucleosomes due to a lack of transcription-coupled histone recycling leads to a disordered chromatin fibre structure, which underlies the spurious aggregation of nearby active genes.

## Discussion

Chromatin states are dynamically disrupted and restored in the wake of replication and transcription to ensure faithful genome regulation within and across cell generations. Here, we show that FACT functions as a general factor for chromatin disruption and restoration in mammalian DNA replication and transcription (Figure 7), reflecting a common mechanistic basis for how DNA-templated processes progress through chromatin. Without FACT, replication forks halt in a stable checkpoint-blind state as they move away from initiation zones and newly synthesized DNA, though limited, fails to be properly assembled into chromatin. Likewise, RNA polymerase elongation rates decline rapidly upon FACT loss, and nucleosome organization collapses in their wake due to a lack of histone recycling. Without retention of pre-existing histones, histone modifications are depleted across TSSs and gene bodies. Transcription-coupled histone recycling, therefore, underpins the chromatin state of active genes. Severe local disruption of chromatin fiber architecture propagates to the 3D genome, leading to aberrant coalescence of active genes into AMCs. This positions nucleosome organization and dynamics as key regulators of spatial genome architecture.

**Figure 7.**
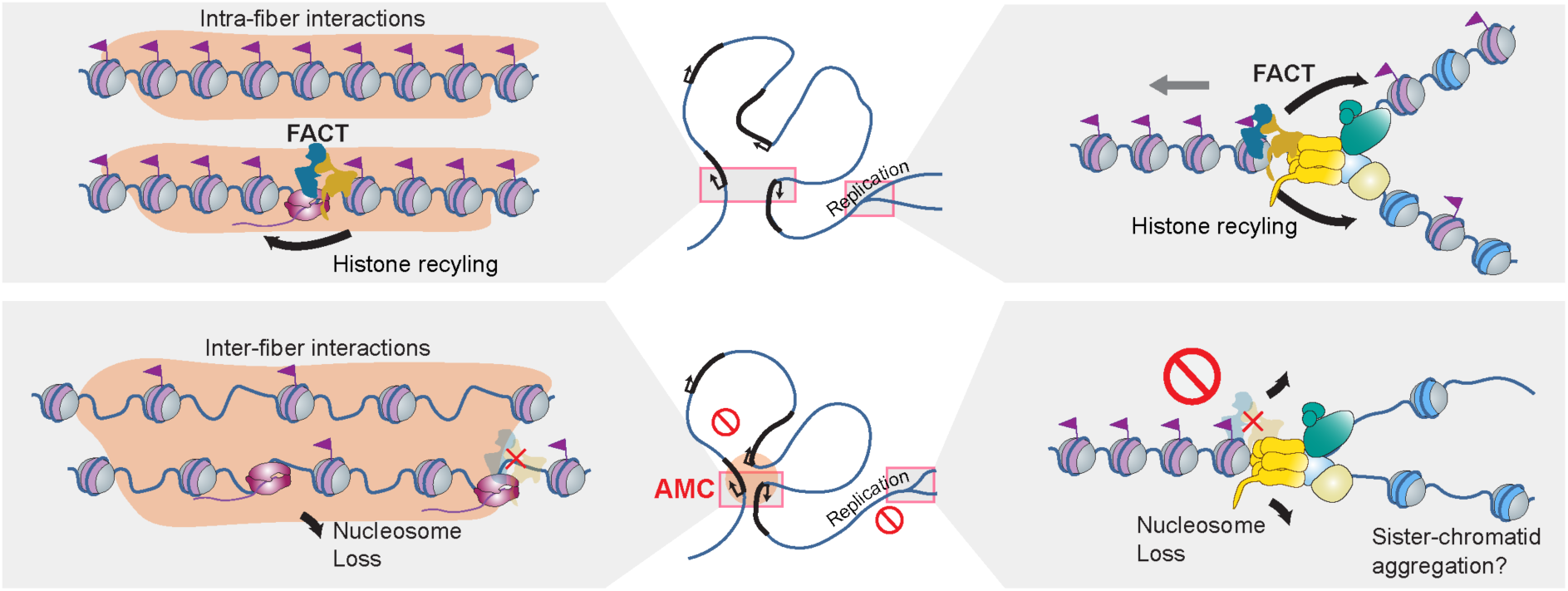
Model. FACT facilitates nucleosome dissolution and restoration during chromatin replication and transcription in mammalian cells. Restoration of well-ordered chromatin fibers by histone recycling counteracts coalescence of active genes into AMCs by favouring intra-fiber interaction over inter-fiber interaction. Likewise, proper nucleosome organization on newly synthesized DNA might reduce the propensity for sister-chromatid aggregation. See discussion for details.

We observe acute arrest of DNA replication and strong global repression of RNA Pol I and II transcription upon SPT16 depletion with loss of viability within 24 hours, consistent with large-scale CRISPR knockout studies identifying SPT16 and SSRP1 as common essential genes across cancer cell lines and in normal fibroblasts (https://depmap.org^48^). Previous studies observed more moderate and variable effects on replication and proliferation upon SPT16 and SSRP1 depletion with slower and/or less efficient depletion approaches^57,58,60,61,90,91^, implying that some cells may adapt to low FACT levels by repurposing other chaperones, as exemplified by LEDGF/HGDF for transcription in differentiated cells^92^. High transcription and proliferation rates in cancer and embryonic stem cells likely make adaptation more challenging, explaining reports of differential FACT requirement in somatic, embryonic and cancer cells^29,58,93^.

Importantly, our model for acute removal of SPT16 in mESCs reconciles the function of FACT as a general mammalian transcription and replication factor with a large body of work from yeast, structural and in vitro studies^27–29,39–46,54,94,95^ This supports a common mechanistic basis coupling nucleosome disruption and histone-recycling in replication and transcription. The dual functionality of FACT in engaging partially unwrapped nucleosomes and co-chaperoning histones together with integrated histone-binding activities would be central to retain and recycle histones: in replication, FACT co-chaperones H3-H4 together with MCM2^44,56,96–98^ and, likewise, yeast Spt6 (which travels with RNAPII) was proposed to co-chaperone histones with FACT during transcription^99^. Whether other factors proposed to substitute for FACT during transcription in differentiated cells would mimic this behaviour and have the capacity to work in replication remains unknown.

Replication forks proceed through chromatin at a rate of 1-2 kb/min in mammalian cells^100^, encountering 5-10 nucleosomes every minute. Without SPT16, DNA synthesis became almost undetectable as forks arrested genome-wide. DNA synthesis at initiation zones was considerably less affected than surrounding regions, indicating that FACT is dispensable for origin firing as observed in vitro^42^. Whether initiation involves alternative means of chromatin disruption remains to be addressed. Outside of initiation zones, replication forks cannot progress without FACT and arrest genome-wide in a checkpoint blind stage without DNA damage. The most parsimonious explanation is that nucleosomes constitute a physical hindrance when FACT cannot aid their disassembly, preventing DNA unwinding by the CMG and thus arresting replication without exposing ssDNA and triggering checkpoint signaling^101^. This model is supported by recent structural analysis proposing that SPT16 engagement with a partially unwrapped nucleosome encountered by a human replisome will trigger nucleosome disruption and enable fork progression^47^, much alike FACT function in transcription^102^. Nascent chromatin fibers, replicated in the absence of FACT, show elevated accessibility with fewer nucleosomes and more intermediate nucleosome assemblies. This partly resembles the consequences of lacking CAF-1^67^, responsible for replication-coupled assembly of nucleosomes from new histones. Several lines of evidence support a function of FACT in histone recycling^56,103^ by co-chaperoning histones together with replication factors like Mcm2^44,96^ and Mrc1^22^ and interacting with Pol alpha^104^. We conclude that FACT is essential for replication-coupled chromatin assembly in mammalian cells, likely operating to assemble nucleosomes from parental histones in parallel to de novo histone deposition by CAF-1. Whether FACT might also contribute to new histone deposition^105^ – on its own or together with CAF-1 – remains open. In mammalian cells, deposition of new histones is required for efficient replication fork progression^106–108^. Without new histones or CAF-1, forks also arrest in a checkpoint blind state with reduced DNA unwinding^106–108^. It is therefore possible that defective chromatin assembly behind the fork also contributes to fork arrest in FACT-depleted cells. We favour the idea^14^ of crosstalk between the histone recycling and the de novo deposition pathways that converge on a common mechanism to fine-tune fork progression with histone supply and demand. FACT-mediated disruption of nucleosomes ahead of the CMG could be the regulatory step, as global replication rates are unaffected in mutants defective at later steps in the recycling pathways towards lagging^15,19^ and leading strand^20,108,109^.

We find that SPT16 is required for RNA Pol I and RNA Pol II progression in mESCs, consistent with a recent study of SSRP1 depletion in leukemic cells^62^. Slow RNA Pol II progression without FACT leaves genic chromatin in disarray^53,55,110,111^. Our data corroborate and expand the critical and conserved role of FACT in transcription-coupled histone H3-H4 and H2A-H2B recycling^24,41,46^. By analyzing individual chromatin fibers in FACT-depleted cells, we establish that histone recycling by FACT is critical to maintain the well-order nucleosome fiber structure across gene bodies, particularly in TSS proximal regions where nucleosome remodellers normally ensure regular phasing. FACT can also play a role in the remodelling process through its interaction with CHD1 during transcription elongation^45,112–114^. Other histone chaperones that accompany RNA Pol II, such as SPT6^115^, cannot compensate for acute FACT loss in mammalian cells. H3.3 incorporation was elevated by about 50% likely as compensatory mechanisms, as observed in yeast, where Spt6 is required for new H3 incorporation in Spt16 mutants^24^. Even so, nucleosome density was substantially reduced after short-term FACT-depletion and the loss of histone modifications in TSS and gene body regions was even more pronounced. This is consistent with work in yeast and Drosophila, showing mislocalization of transcription-associated modifications when FACT is impaired over multiple cell generations^53,55,110^. In yeast, but not in Drosophila, this was associated with nucleosome loss. Our findings imply that new H3.3-H4 cannot restore the levels of active modifications despite the presence of RNA Pol II, which can recruit histone-modifying enzymes^116^. Collectively, this argues that recycling of pre-existing histones with their modifications is key to maintaining high levels of active modifications in genic regions. This could reflect that turnover exceeds the rates of histone-modifying enzymes and/or a direct coupling of histone recycling to co-transcriptional histone modification as recently described for SETD2-mediated H3K36me3 deposition^117,118^.

Disorder at the nucleosome level as a consequence of FACT depletion rapidly alters 3D genome organization. At least two processes drive genome folding: active loop extrusion and affinity-driven genome compartmentalization^119,120^. Compartments are suggested to form by block copolymer phase separation, primarily driven by affinities between B compartments^121^. However, little is known about the mechanisms behind the coalescence of genomic regions into global chromatin domains, in particular A compartments. Micro-compartments (MCs) have been observed with high-resolution capture strategies between distal regulatory elements and transcription start sites^75^. Recently, MCs were shown to intermediately form during mitotic exit and their prevalence can be extended by inhibiting the import of chromatin factors, such as the loop extrusion and transcription machinery, into the newly formed nucleus^76,77^. The AMCs found in our system are distinct in two ways from previously identified MCs. First, upon perturbation, AMCs form rapidly in interphase cells, excluding cell cycle effects. Second, compared to previously identified MCs that predominantly form between genes and enhancers, AMCs appear to be restricted to genes. However, AMCs can form in trans like compartments and MCs. Loop extrusion antagonizes the coalescence of block copolymers in simulations^120,122^, and impaired loop extrusion strengthens genome compartments^83–85,123^. In yeast, SPT16 was suggested to maintain Cohesin stability on chromatin^124^. However, in FACT-depleted cells, Cohesin is not lost, loop-extrusion-dependent CTCF-CTCF interactions are only moderately weakened, and Cohesin occupancy is not a good predictor of AMC formation. AMCs thus form in a loop-extrusion independent process, but it is possible that a moderate reduction in loop extrusion across genes (linked to RNA Pol II accumulation^125^) might contribute by reducing ‘untangling’ of AMCs.

Using machine learning, we found that high expression along with low nucleosome and H2A.Z occupancy after SPT16 depletion are the most predictive features that separate AMC-forming and non-forming genes. Nucleosome organization is more severely perturbed at AMC-forming genes, displaying reduced nucleosome occupancy and diffuse nucleosome repeat lengths with a tendency towards larger nucleosome spacing. The effect of mesoscale chromatin structure, such as nucleosome fibers, on genome compartmentalization is poorly understood. Evidence from biochemical chromatin reconstitution experiments supports the idea that the nucleosome fiber structure and chromatin remodelers can regulate the ability of chromatin to coalesce^126–128^, with NRL and nucleosome occupancy regulating the balance between intra-fiber and inter-fiber contacts. Our work establishes that nucleosome fiber structure – and the levels of disorder – impact chromatin aggregation in cells. We demonstrate that proper chromatin restoration of nucleosome organization by FACT prevents spurious aggregation of nearby active genes. This argues that genome compartmentalization is dynamic and can be rapidly modulated by altering chromatin composition. Recent ultra-deep Hi-C analysis revealed that compartment organization is generally more fine-scaled than first anticipated with an average size of 12.5 kbp, and including compartment type interaction at the beginning of active genes^129^. We envision that such active genes, in spatial proximity, coalesce into strong compartments upon loss of ordered chromatin fiber structure, because it shifts the propensity from intra-fiber to inter-fiber interactions (Figure 7). Contributing forces are likely multi-valent interactions of chromatin and transcriptional regulators binding the TSS regions along with somewhat reduced loop-extrusion across the affected genes. We note that hyperacetylation leads to increased interactions between chromatin domains in cells, likely explained by acetylation-binding proteins with propensity to coalesce, such as BRD4^130^. Importantly, our data show that - in addition to transcriptional regulation^131^ - the highly ordered nucleosome structure in TSS proximal regions serves a role in preventing spurious gene aggregation. It is attractive to envision that perturbation of nucleosome structure at other locations, such as sister-chromatids in FACT depleted cells, may similarly impact self-association of distinct chromatin environments.

## Limitations of the study

The consequences of SPT16 depletion were addressed in mESCs, and it is possible that other chaperones substitute for FACT in other cell types due to altered expression levels, as evidenced for LEDGF/HGDF2 in myoblasts^92^. However, the widespread dependence of mESCs on FACT allows us to obtain a comprehensive understanding of the functions of the mammalian protein complex, as compensatory effects do not mask the phenotypic changes. Due to the severe effect of FACT depletion on replication activity and EdU incorporation, we were unable to assess the effects on the histone modification landscape of newly replicated chromatin. In the future, FACT separation of function mutants (disabling histone recycling and not nucleosome disruption) might make this possible. Similarly, the strong impairment of RNA Pol II progression and rapid loss of cell viability precluded a deeper investigation into the downstream transcriptional effects of gene aggregation. Other models will thus be required to understand the physiological importance of preventing extensive inter-fibre contacts between neighbouring gene bodies.

## Resource Availability

### Lead contact

Further information and requests for resources and reagents should be directed to and will be fulfilled by the lead contact, Anja Groth.

## Acknowledgements

We thank Jesper Svejstrup and members of our teams for fruitful discussions, Smaragda Kompocholi and Lea Gregersen for training in TTchem-seq, and the staff of the CPR/reNEW Genomics Platform, the CPR Protein Imaging Platform, and the ReNEW Flow Cytometry platform for support. Research in N.K.’s laboratory was supported by the Lundbeck Foundation (R368-2021-1076). Research in A.v.O.’s laboratory was supported by the European Research Council (ERC-AdG no. 101053581). Research in A.G.’s laboratory was supported by the Danish National Research Foundation (DNRF195), the Novo Nordisk Foundation (NNF21OC0067425), the European Research Council (ERC-AdG no.101142230), and Independent Research Fund Denmark (DFF3101-00149B). Q.D. is a NHMRC Investigator grant recipient (1177792). Research at Novo Nordisk Foundation Center for Protein Research is supported by the Novo Nordisk Foundation (NNF14CC0001, NNF24SA0098829).

## Author Contributions

V.F., R.C.M.F., N.K. and A.G. conceived the study and R.C.M.F., R.R.J., V.F., J.v.d.B, A.B., N.K., and A.G. designed experiments. A.G., N.K. V. R. and A.v.O. supervised the study. V.F, R.C.M.F. and A.B. performed cell line validation, flow cytometry, and EdU-seq. R.C.M.F and A.B performed microscopy and viability assays. R.C.M.F analysed EdU-seq with assistance from V.F. and Q.D. J.v.d.B. performed scEdU-seq and analysed scEdU-seq data. R.C.M.F. performed TTchem-seq and analysed data with assistance from N.A.. R.C.M.F and S.W performed RASAM and R.C.M.F analysed data with assistance from H.R. and M.Y.. R.C.M.F performed ChIP-seq and analysed data with assistance from J.O.L.. R.R.J performed Micro-C and J.O.L. and N.K. analysed data. R.R.J performed Hi-C and N.K analysed the data. R.C.M.F, R.R.J, J.O.L., N.K., and A.G. wrote the manuscript with input from all authors.

## Methods

### Cell culture

E14 mESCs were grown on feeder-free plates coated with 0.2% gelatin in DMEM supplemented with 15% FBS (Invitrogen), 1x penicillin/streptomycin, 1x non-essential amino acids, 1x β-mercaptoethanol, and custom-made LIF at 37C and 5% CO_2_. Cells were passaged every 48 hours using TrypLE and regularly tested for mycoplasma. Biological replicates were cultured separately for a minimum of 2 passages prior to measurement. dTAG treatments were performed with 1 μM dTAG or an equal volume of DMSO as a vehicle control for the specified time prior to labelling and/or harvesting.

Drosophila S2-DRSC cells were obtained from the Drosophila Genomics Research Centre. S2 cells were grown in suspension in spinners in M3+BPYE media: Shields and Sang M3 Insect Medium (Sigma, S-8398), KHCO3 (Sigma, 12602), yeast extract (Sigma, Y-1000), bactopeptone (BD, 211705), 10% heat-inactivated FBS (GE Hyclone, SV30160.03) and 1X penicillin/streptomycin (GIBCO, 151400122). Cells were incubated at 25°C with 5% CO_2_.

### Cloning and Cell Line Generation

Homology arms corresponding to the C-terminus of Supt16h were amplified from E14 Ju mESC genomic DNA using primer sequences in Supplementary Table 2 and integrated into a pUC19 plasmid by TOPO-TA cloning using the TOPO TA cloning kit (Invitrogen, 11543147). The dTAG construct was obtained from a BAP1-linker-dTAG donor plasmid (gift from the Klose lab^19^) and was inserted into the pUC19 with homology arms by megaprimer cloning ahead of the Supt16h stop codon. The gRNA plasmid was generated from the px459 plasmid (Addgene #62988) as described in^132^ by introducing the gRNA sequence AGAGAAAGTAATATGAACCT targeting the C-terminus of Supt16h. To generate the cell line, 5x10^5^ E14 Ju mESCs were seeded on a 6-well plate 24 hours prior to transfection. Cells were transfected with 0.5 μg of gRNA plasmid and 2 μg of dTAG donor plasmid using Lipofectamine 3000 (Invitrogen L3000001) according to manufacturer instructions. Transfected cells were seeded sparsely 24 hours later on a 10 cm dish and selected with puromycin (2 μg/ml) for 48 hours. Surviving colonies originating from a single cell were picked and expanded and the clones validated by genotyping and Sanger sequencing. Two clonal cell lines were used for further experiments and were validated for similar viability, replication activity, and transcription levels.

### Western Blotting

Cells were seeded on 6-well plates 48 hours before collection. Treatment with DMSO or 1 μM dTAG-13 was performed immediately before harvesting. To harvest, cells were washed twice with PBS and Laemmli sample buffer (60 mM Tris-HCl pH 6.8, 2% SDS, 10% glycerol, 5% β-mercaptoethanol, 1% bromophenol blue) was added directly to the well. Lysates were collected, 25U Benzonase (Sigma) added and incubated for 1 hour at 37°C before being boiled for 10 minutes at 95°C. SDS-PAGE was performed on a 4-12% Bis-Tris gel and proteins transferred to a nitrocellulose membrane by semi-dry transfer. Memcode staining was performed to assess protein transfer. Membranes were blocked in 5% milk in PBST and incubated overnight at 4°C with primary antibodies diluted in 5% milk. Membranes were washed 3x in PBST then incubated for 1 hour at room temperature with HRP-conjugated secondary antibody (1:15000), followed by 3x washes in PBST. Signal was detected with Pierce ECL or ECL Pico Plus Western Blotting Substrate (ThermoScientific) with a ImageQuant LAS 4000 Camera (GE Healthcare).

### CellTiter Blue viability assay

3x10^4^ cells were seeded 48 hours before CellTiter Blue (Promega G8081) reagent addition in 96-well plates with 6x technical replicates per condition. 3x empty wells with 100 μl of medium were used as a blank control. dTAG or DMSO treatment was performed for the specified time before addition of CellTiter Blue reagent. 20 µl CellTiter Blue reagent was added to 100 µl of medium and incubated for 4 hours prior to measurement. Fluorescence in each well was measured using an Omega Fluostar. Viability percentages were calculated by comparing fluorescence in the dTAG condition with the time-matched DMSO control for each time point.

### High-throughput microscopy

3x10^5^ mESCs were seeded on laminin-coated 96-well plates and cultured overnight in standard conditions. dTAG or DMSO treatment was started by changing medium to treatment-containing medium. For labelling ongoing replication, cells were incubated with 10 μM EdU (Invitrogen, 11590926) for 10 minutes, EdU medium was removed and ice-cold PBS used to wash the wells. 5-ethynyl uridine (5-EU, Invitrogen E10345) labelling was performed with 100 mM 5-EU for 30 minutes, after which ice-cold PBS was used to wash the cells prior to fixation. Cells were fixed in freshly prepared 4% formaldehyde in PBS for 15 minutes at room temperature. Formaldehyde was removed by washing 3x in PBS and cells were permeabilised for 20 minutes in 0.1% triton-X in PBS. Labelled DNA/RNA was visualised using the Click-IT EdU Alexa Fluor 647 kit (Invitrogen C10643) according to manufacturer instructions. After the Click-IT reaction, cells were blocked in blocking buffer (0.1% Triton, 3% BSA, 10% FBS in PBS) for 1 hour at RT. For plates with antibody staining, antibody incubations were performed overnight at 4°C with the following concentrations (gH2A.X 1:1000, Chk1-Ser317P 1:1000). Plates were washed 3x in blocking buffer and incubated with secondary antibody (anti-Rabbit Alexa 568, anti-Mouse Alexa 488, both 1:1000) in blocking buffer for 1 hour at RT. Plates were then washed 2x in blocking buffer and 3x in PBS, with DAPI added during the second wash. Each well was kept in 100 μl of PBS for imaging. Imaging was performed on an Opera Phenix microscope at 20x magnification. Nuclei segmentation using DAPI and mean intensity calculations for individual nuclei were performed with Harmony 4.2 software and RStudio was used for further analysis and visualisation.

### Flow Cytometry

2.4 x10^5^ mESCs were seeded 24 hours prior to harvesting on a 6-well plate. Cells were treated with 1 μM dTAG-13 or volume-matched DMSO for the specified times. After treatment, cells were labelled for 10 minutes with 10 μM EdU. Cells were detached with TrypLE and pelleted by centrifugation at 300g for 5 minutes. Cells were fixed in 70% EtOH at -20°C and stored until further analysis.

Stored cells were thawed and pelleted by centrifugation at 300g for 4 minutes. Cells were washed 2x in 1% BSA-PBS and permeabilised with 0.25% Triton-X in PBS for 10 minutes at room temperature. Cells were washed once in 1% BSA-PBS and EdU click reaction was performed by incubation in click reaction buffer (Invitrogen, C10419) (2 mM CuSO_4_, 1:200 Alexa Fluor 647-Azide, 1x Click-iT Buffer Additive in PBS) for 35 minutes at room temperature in the dark. Cells were washed once with 1% BSA-PBS and incubated with 10 μg/μl proprium iodide and 20 μg/μl RNAse A for 30 minutes at room temperature. Cells were kept at 4 °C until analysis on an LSR Fortessa flow cytometer.

### EdU-seq

4x10^6^ mESCs were seeded 48 hours prior to collection on a 15cm plate. 1 μM dTAG or equal volume DMSO treatment was added for the specified time before EdU labelling. 10 μM EdU was added to each sample for 10min prior to harvesting. To harvest, medium was removed and plates were washed 2x with ice-cold PBS before incubation for 5 minutes in 1% formaldehyde to halt replication. Plates were then washed 2x with PBS and cells scraped and transferred to a 50ml tube in PBS. Cell pellets were centrifuged for 5 minutes at 1000g, PBS removed, and the cell pellets flash frozen in liquid nitrogen. Pellets were stored at -80°C until further use.

Nuclei extraction was performed according to^133^. Chromatin sonication was validated using a TapeStation and total chromatin was quantified by Qubit. EdU-seq libraries were prepared with 0.05% spike-in EdU labelled S2 chromatin. For EdU-seq the protocol was performed as described in the ChOR-seq protocol^133^, omitting the chromatin immunoprecipitation steps. 100 μl DNA in TE buffer was decrosslinked overnight by incubation for 30 minutes with 5 μl of RNase A (10mg/ml) at 37°C followed by addition of 1 μl proteinase K (10mg/ml), 2.5 μl of 20% SDS and 2 μl of 5M NaCl and incubation for 10 hours at 37°C followed by 6 hours at 65°C. DNA was then purified by MinElute and quantified by Qubit. Sequencing adapters were ligated to DNA according to kit instructions and purified using AMPure XP beads and eluted in water. Biotin was conjugated to EdU-labelled DNA by incubation by making the sample up to 100 μl volume of Click buffer (final concentration: 1x PBS, 0.5 mM picolyl-azide-PEG4-biotin, 0.5 mM THPTA, 0.1 mM CuSO_4_, and 10 mM sodium ascorbate). Click reaction was cleaned by AMPure XP beads, and samples eluted in 20 μl of water. 10 μl MyOne T1 streptavidin Dynabeads per sample were washed 3x with 1x BWT buffer (10 mM Tris-HCl pH 7.5, 1 mM EDTA, 2 M NaCl, 0.1% Tween-20), before being resuspended in 20 μl 2x BWT buffer per sample. 20 μl of streptavidin beads in 2x BWT buffer was added to each sample, and biotinylated DNA captured by incubating with rotation for 30 minutes at 37°C. Beads were washed 4x with 1x BWT buffer and DNA eluted in 0.1x TE buffer before library preparation. Libraries were prepared with KAPA hyperprep kit (Roche, 07962363001) according to manufacturer instructions and sequenced single-end on a NextSeq 2000.

### scEdU-seq

1x10^6^ mESCs were seeded on a 10cm plate 48 hours prior to harvesting. Cells were labelled with 10 μM EdU for 15 minutes prior to dTAG or DMSO treatment. EdU was washed out by 4x washes in full medium, and replaced with medium containing 1 μM dTAG or the same volume of DMSO. After 1 hour of incubation, the second EdU treatment was performed for 15 minutes with 10 μM EdU. Cells were detached from the plate using TrypLE and washed 2x in PBS. Cells were resuspended in 1 ml of ice-cold PBS and 3 ml of ice-cold 100% (f.c. 80%) ethanol was added for fixation overnight at -20C. After fixation, cells were pelleted then resuspended in storage buffer (10% DMSO, 20 mM HEPES pH 7.5, 150 mM NaCl, 24.768 μM spermidine, 0.05% Tween-20, 2 mM EDTA pH 8) with protease inhibitors. Resuspended cells were stored at -20C until further analysis.

scEdU-seq was performed as previously described^65^. Briefly, mESCs were resuspended and washed in 1 mL of Wash Buffer (20 mM HEPES, pH 7.5; 150 mM NaCl; 25 mM spermidine; 0.05% Tween; and 2 mM EDTA). Biotin-PEG3-Azide (10 mM, Sigma-Aldrich) was then conjugated to the EdU molecules using a CuAAC click reaction. This was followed by staining with DAPI (10 μg/mL) in PBS containing 0.25% BSA. mESCs were sorted into 384-well plates for scEdU-seq processing. Library preparation involved several steps: proteinase K digestion, genome digestion with NlaIII, DNA blunt-ending, A-tailing, and adapter ligation with incorporated cell barcodes and unique molecular identifiers (UMIs). The resulting single-cell libraries were pooled and immobilized on Dynabeads MyOne Streptavidin C1 (Thermo Fisher Scientific) to capture DNA replication fragments. These fragments were subsequently released by heat denaturation and filled in using the Klenow enzyme. Amplification of the libraries was performed through in vitro transcription (IVT), reverse transcription (RT), and PCR, followed by sequencing on an Illumina NextSeq 1000 platform (P3, 2×100 bp). The code for analysis and plotting is available on GitHub: https://github.com/jervdberg/XXXX

### RASAM

Replication-aware single-molecule accessibility mapping (RASAM) was performed as in^67^. In brief, SPT16-dTAG mESCs were incubated with 0.5 μM dTAG-V1 or volume-matched DMSO for 2 hours prior to labelling. Medium was then replaced with medium containing dTAG or DMSO with 10 μM BrdU (abcam ab142567) cells were incubated in BrdU-containing media for 1 hour. After labeling, samples were rinsed three times with 1x PBS and collected using TrypLE Express Enzyme (1X) (12605010). all nuclei were collected by centrifugation (300xg, 5 min), washed in ice cold 1X PBS, spun again (300xg, 5 min), and resuspended in 1 mL Nuclear Isolation Buffer (20mM HEPES, 10mM KCl, 1mM MgCl2, 0.1% Triton X-100, 20% Glycerol, and 1X Protease Inhibitor (Roche 4693132001)) per 10 million cells by gently pipetting 5x with a wide-bore pipette tip. The suspension was incubated on ice for 5 minutes, and nuclei were pelleted (600xg, 4°C, 5 min), washed with Buffer M (15 mM Tris-HCl pH 8.0, 15 mM NaCl, 60 mM KCl, 0.5 mM Spermidine), and pelleted by centrifugation (600xg, 4°C, 5 min).

Nuclei were resuspended in 200 μL of Methylation Reaction Buffer (Buffer M containing 1mM S-adenosylmethionine (SAM)) (New England BioLabs B9003S). 20 μL high-concentration EcoGII was added per 1x10^6^ nuclei and the nuclei suspension was incubated at 37°C for 30 minutes. An additional 1 μL of 32 mM SAM was supplemented after 15 minutes.

After the EcoGII incubation, 2.65 μL 10% SDS and 2.65 μL of 20 mg/mL Proteinase K (Thermo Scientific AM2548) were added and incubated at 65oC for 2 hours. To extract the DNA, an equal volume of Phenol-Chloroform was added and mixed by shaking. Samples were centrifuged at max speed for 2 min and the aqueous portion removed. 0.1x volumes of 3M NaOAc, 1 μL GlycoBlue, and 3x volumes of cold 100% EtOH were added, mixed by inversion and incubated overnight at -20°C. Samples were then centrifuged at max speed for 30 minutes at 4°C, washed with 500 μL 70% EtOH, and centrifuged at max speed for 2 minutes at 4°C. The resulting pellet was air dried and resuspended in 40 μL Buffer EB. Sample concentrations were measured via Qubit High Sensitivity DNA Assay.

Purified DNA was sheared using a Covaris g-tube (520079) in a 5424 rotor at 7,000 RPM for 6 passes for a target size between 6,000 and 8,000 bp. Sheared DNA was used as input for the PacBio HiFi SMRTbell Library Preparation protocol using the SMRTbell Express Template Prep Kit 2.0. Briefly, samples underwent removal of single stranded overhangs, DNA damage repair, end-repair, A-tailing, barcoded SMRTbell adapter ligation, and exonuclease cleanup according to manufacturer instructions, followed by a 1X AMPure PB bead cleanup. Final sample concentration was measured via Qubit High Sensitivity DNA Assay and library size was measured on an Agilent Bioanalyzer DNA chip. Libraries were sequenced on PacBio Sequel II 8M SMRTcells.

### TTchem-seq

TTchem-seq was carried out as described in^66^. 1.2x10^6^ mESCs were seeded 48 hours before harvesting, leading to a final cell number of around 10x10^6^. dTAG treatment was performed for the specified time prior to RNA labelling. RNA was labelled in vivo with 1 mM 4-SU (Glentham Life Sciences, GN6085) for 15 minutes prior to the addition of TRIzol (Thermo Fisher Scientific). RNA was extracted by TRIzol according to manufacturer instructions and quantified by Qubit BR RNA assay.

100 μg of mouse RNA was mixed with 1 μg of 4TU-labelled yeast RNA and brought to a total volume of 100 μl with water. The mix was fragmented by adding 20 μl 1 M NaOH and incubating on ice for 20 min. 80 μl of 1 M Tris-HCl pH 6.8 was added to stop fragmentation and samples were cleaned up twice with Micro Bio-Spin P30 Gel Columns (BioRad, 7326250) following manufacturer instructions. Biotinylation of 4SU and 4TU residues was carried out in a total volume of 250 μl 10 mM Tris-HCl pH 7.4 and 1 mM EDTA, containing MTSEA biotin-XXlinker (Biotium, BT90066) for 30 min at room temperature in the dark. RNA was then purified by phenol:chloroform extraction, denatured for 10 min at 65°C and added to 200 μl μMACS Streptavidine MicroBeads (Miltenyi Biotec, 130-074-101). After 30 min incubation at room temperature, the mix was loaded to a μColumn in the magnetic field of a μMACS magnetic separator. Beads were washed twice in a 55C buffer containing 100 mM Tris-HCl pH7.4, 10 mM EDTA, 1M NaCl and 0.1% Tween20. Biotinylated RNA was eluted twice in 100 mM DTT (200 μl final volume) and cleaned up with the RNeasy MinElute kit (QIAGEN, 74204) using 1050 μl 100% ethanol and 750 μl RLT buffer to precipitate RNA <200nt.

TTchem-seq libraries were prepared from 5ng of purified RNA using the NEBNext® Ultra™ II Directional RNA Library Prep Kit for Illumina® following the protocol for FFPE RNA with adapters added from the NEBNext® Multiplex Oligos for Illumina (Unique Dual Index UMI Adaptors RNA Set 1) according to manufacturer instructions. Average fragment size of the final library was determined on a TapeStation using the HS 5000 DNA kit, and concentrations determined by Qubit. Sequencing was performed single-end on a NextSeq 2000.

### ChIP-seq

4x10^6^ mESCs were seeded 48 hours prior to collection on a 15cm plate. 1 μM dTAG or equal volume DMSO treatment was added for the specified time before harvesting. To harvest, medium was removed and plates were washed 2x with PBS before incubation for 10 minutes in 1% formaldehyde. Plates were then washed 2x with PBS and cells scraped and transferred to a 50 ml tube in PBS. Cell pellets were centrifuged for 5 minutes at 1000g, PBS removed, and the cell pellets flash frozen in liquid nitrogen. Pellets were stored at -80°C until further use.

Nuclei extraction was performed according to^133^, and sonication was performed on a Covaris E220 Evolution with 10% duty factor. Chromatin sonication was validated using a TapeStation and total chromatin was quantified by Qubit. ChIP-seq spike-in was prepared from Drosophila S2 chromatin and added to 1% of the final chromatin input for each ChIP. RNA Pol II, RAD21, RING1B, and SUZ12 ChIPs were performed with 50 μg, H2BUb and H2A.Z with 25 μg, and H3K4me3, H3K36me3, H3K27ac, H3.3, H3, and H2B ChIPs were performed with 10 μg of input chromatin. Chromatin was incubated with antibodies overnight at 4°C before incubation with Protein A/G beads for 3 hours at 4°C. For histone ChIPs, beads were rinsed 2x with low-salt (150 mM) RIPA buffer, washed 1x with low-salt RIPA buffer for 5min, washed 3x with high-salt (500 mM) RIPA, and 1x in LiCl wash buffer before being eluted in 1x TE. For non-histone ChIPs, 2x low-salt rinses, 2x low-salt washes, 1x high-salt wash, and 1x LiCl wash were performed. ChIP DNA was quantified by Qubit and KAPA Hyperprep kit was used for library preparation. Samples were sequenced paired-end with 56bp reads on a NextSeq 2000.

### Micro-C

#### Crosslinking

Cells were harvested as previously described^74^ and resuspended in 1x DPBS w/o Ca^2+^ and Mg^2+^ to reach a solution of 1x10^6^ cells/ml. The cell suspensions were then added 1% Formaldehyde and incubated at RT for 10 minutes on a rotator. The reaction was quenched using 0.25 M glycine and incubated at RT for 5 minutes. The samples were centrifuged at 1000g for 5 minutes, washed once with 1x PBS w/o Ca^2+^ and Mg^2+^ with 0.1% BSA and resuspended in 1x PBS w/o Ca^2+^ and Mg^2+^ with 0.1% BSA, reaching a solution of 4x10^6^ cells/ml. EGS were dissolved in DMSO and 3 mM EGS were added to the cell suspension prior to incubation at RT for 40 minutes on a rotator. The reaction was quenched using 0.4 M glycine and incubated at RT for 5 minutes. The cells were centrifuged 1000g for 5 minutes and resuspended in 1x PBS w/o Ca^2+^ and Mg^2+^ reaching a concentration of 5x10^6^ cells/ml. The crosslinked cells were aliquoted into Eppendorf tubes to generate samples either containing 5x10^6^ or 1x10^6^ cells. The aliquoted samples were centrifuged 2000g for 5 minutes and supernatant were carefully removed. Pelleted cells were snap frozen in liquid nitrogen and stored at -80°C until further processing.

#### MNase titration

For all conditions in each experimental setup, the optimal MNase digestion was estimated by MNase titration. One vial of 1x10^6^ cells were thawed on ice for 10 minutes, resuspended in 0.5ml 1x PBS w/o Ca^2+^ and Mg^2+^ with 0.1% BSA and incubated on ice for 20 minutes. The cells were centrifuged at 10000g for 5 minutes and resuspended in 0.5 ml MB1 buffer (50 mM NaCl, 10 mM Tris-HCl pH 7.5, 5 mM MgCl_2_, 1 mM CaCl^2^, 0.2% NP-40, 1x Roche cOmplete EDTA-free Protease Inhibitor) to extract the nuclei. Samples were centrifuged at 10000g for 5 minutes and pelleted nuclei were resuspended in 0.2 ml MB1 buffer and distributed equally into four tubes. MNase was diluted into four different concentrations: 10 U, 2.5 U, 0.625 U and 0.1256 U and 1 µl from each dilution were added to one tube each in a 15-second interval and incubated at 37°C for 10 minutes. With the same order and 15-second interval, 200 µl STOP-buffer (150 µl 10 mM Tris-HCl, 25 µl 10% SDS, 25 µl 20 mg/ml Protein Kinase, 2 µl 0.5 M EGTA) was added to each tube and incubated at 65°C for 2 hours on a shaker. 0.5 ml Phenol-Chloroform-Isoamyl-Alcohol was added to each sample, samples mixed thoroughly by vortexing and centrifuged at 19800g for 5 minutes. The upper aqueous phase containing the DNA was transferred to new Eppendorf tubes and purified using the Qiagen PCR purification kit eluted in 12 µl. All samples were run on a 1.5% agarose gel and the proper MNase concentration were estimated from the resultant MNase digestion ladder.

#### Micro-C Preparative libraries

To prepare the Micro-C libraries, one vial of 5x10^6^ cells were thawed on ice for 10 minutes and resuspended in 1 ml 1x PBS w/o Ca^2+^ and Mg^2+^ with 0.1% BSA and incubated on ice for 20 minutes. Cells were centrifuged 10000g for 5 minutes and resuspended in 0.5 ml MB1 buffer. Samples were centrifuged at 10000g, and pelleted cells were resuspended in 1 ml MB1 buffer and distributed equally in 5 new Eppendorf tubes with 200 μl/tube. With a 15-second interval, 1 µl MNase with the estimated concentration from the MNase titration was added each tube and tubes incubated at 37°C for 10 minutes. With the same order and same 15-seconds interval, 1.6 µl EGTA was added to each tube and tubes incubated at 65°C for 10 minutes. Samples were centrifuged at 10000g, and the pelleted nuclei were resuspended in 0.5 ml 1x ligase buffer pooling the five tubes into one. Samples were centrifuged at 10000g and resuspended in 0.5 ml 1x ligase buffer. 10% of each suspension were transferred to clean Eppendorf tubes, added 150 µl Tris-HCl, 25 µl 10% SDS and 25 µl 20 ng/ml Proteinase K and incubated overnight at 65°C. The remaining samples were centrifuged at 10000g, and the pellets were resuspended in 90 ul/sample End-Chewing master mix (10x ligase buffer, H2O, 10 U/µl T4 PNK) and incubated at 37°C for 15 minutes with mixing. Following incubation, 10µl Klenow Fragment was added to each sample and samples incubated at 37°C for 15 minutes. 50 µl End-labeling master mix was then added to each sample (1 mM Biotin-dATP, 1 mM Biotin-dCTP, 10 mM dTTP+dGTP, 10x ligase buffer, 20 mg/ml BSA and H_2_O) and incubated at 25°C for 45 minutes with mixing. To the samples was added 9 µl 0.5 M EDTA and samples incubated at 65°C for 20 minutes, followed by centrifugation at 10000g for 5 minutes. Pelleted nuclei were washed once in 0.5 ml cold 1x ligase buffer, and then resuspended in 500 µl Ligation master mix (10x ligase buffer, 20 mg/ml BSA, 400 U/µl T4 DNA ligase and H_2_O) and incubated at room temperature for 2.5 hours. Nuclei were then pelleted at 10000g for 5 minutes and resuspended in 0.2 ml Exonuclease master mix (10x NEB buffer 1, 100 U/l Exonuclease III and H_2_O) and incubated at 37°C for 15 minutes. Samples were then added 25 µl 10% SDS and 25 µl 20 mg/ml Proteinase K and incubated overnight at 65°C. All samples were added 0.5 ml Phenol-Chloroform and centrifuged at 19800g for 5 minutes and the aqueous phase was transferred to new Eppendorf tubes. The DNA was purified using the Qiagen PCR purification kit eluting input samples in 15 µl EB and library samples in 30 µl EB. The samples were added loading buffer and run on a 1.5% agarose gel at 120 V until bands were properly separated, comparing input samples and library samples as a quality control for the ligation efficiency. For all the library samples, the 350 bp band was cut out of the gel and purified using the Qiagen gel purification kit eluting the samples in 150 µl EB.

### On-bead library prep

To bind the DNA onto the streptavidin beads, 10 µl beads per sample were washed twice with 0.5 µl TBW buffer (5 mM Tris-HCl pH 7.5, 0.5 mM EDTA, 1 M NaCl, 0.05% Tween20) and the washed beads were resuspended in 150 µl per sample 2x B&W buffer (10 mM Tris-HCl pH 7.5, 1 mM EDTA, 2 M NaCl). Each library sample were then added 150 µl of the beads-solution (1:1) and incubated with rotation at room temperature for 20 minutes. The beads now containing the DNA fragments were placed in a magnet and the cleared supernatant were removed. The beads were then washed twice in 300 µl TBW buffer and washed once in 100 µl 0.1% TE buffer before being resuspended in 50 µl 0.1% TE buffer and transferred to clean PCR-tubes. The samples were then processed using the NEBNext® Ultra^TM^ II DNA Library Prep Kit for Illumina® protocol step 1 and 2, following vendors instructions. Following end-prep and adapter ligation, the tubes were placed in a magnet and the supernatant was removed. The beads were washed once with 200 µl TBW and once with 100 µl 0.1% TE buffer. The beads were then resuspended in 20 µl 0.1% TE buffer and 1 µl from each sample were amplified in a 10 µl PCR reaction with 16 cycles to estimate the minimal number of cycles for the library PCR. Subsequently, the main samples were amplified in a 200 µl reaction with the appropriate number of cycles estimated from the 1 µl PCR test, and the amplified samples were purified with SPRI Select AMPure beads in the ratio 1:0.9 to get rid of potential adapter dimers. The AMPure purified samples were eluted in 20 µl 0.1% TE buffer and were measured by Qubit. All samples were run on the TapeStation to estimate the average fragment size and diluted to 4 nM pools for paired-end sequencing. Samples were sequenced on the NextSeq2000 or NovaSeq.

### Hi-C 3.0

Hi-C 3.0 was performed as previously described^134^, with minor modifications. 5x10^6^ double-crosslinked cells were lysed in lysis buffer (10 mM Tris-HCl pH 8.0, 10 mM NaCl, 0.2% Igepal, 1x Roche cOmplete EDTA-free Protease Inhibitor) and digested for >16 hours with 10 U/µl Ddel and 50 U/µl DpnII, before incubating the samples for 4 hours with Biotin-fill-in-master mix (7 µl 10x NEBuffer 3.1, 1.5 µl 10 mM dCTP, 1.5 µl 10 mM dGTP, 1.5 µl 10 mM dTTP, 14 µl 1 mM biotin-dATP, 10 µl 5U/µl Klenow DNA polymerase, H_2_O). The proximal DNA fragments were ligated by adding 665 µl ligase master mix (120 µl 10x ligation buffer, 120 µl 10% Triton X-100, 12 µl 10 mg/ml BSA, 33.25 µl 400 U/µl T4 DNA ligase, H_2_O) to each sample and samples were incubated over night at 16°C with shaking. 50 µl 20 mg/ml proteinase K was added to each sample and incubated overnight at 65°C with shaking. For each sample, the DNA was purified with 2 volumes of Phenol-Chloroform-Isoamyl-Alcohol and the aqueous phase was transferred to clean tubes. The aqueous phase from each sample was then purified by ethanol precipitation and the solubilized pellet resuspended in 450 µl TLE buffer (10 mM Tris-HCl pH 8.0, 0.1 mM EDTA) prior to size selection through a 10 kDa Amicon Ultra-0.5 centrifugal Filter Unit. The remaining samples were eluted in 80 ul TLE buffer and 10 µg DNA from each sample was used for further processing. Biotin-removal master mix (13 µl 10x NEBuffer 2.1, 3.25 µl 1 mM dATP, 3.25 µl 1 mM dGTP, 13 µl 300 U/ml T4 DNA polymerase, H_2_O) was added to each 10 µg DNA sample and incubated overnight according to the protocol. Sample were sonicated using a Covaris E220 according to manufacturer instructions to achieve fragment size distributions <500 bp followed by size selection using SPRI-Select beads to obtain Upper – and Lower fractions. Both fractions were eluted from the beads using 150 µl TE buffer (10 mM Tris-HCl pH 8.0, 1 mM EDTA) and further processed as previously described for Micro-C in the “on-beads-library-prep”-section.

### Sequencing Data Analysis

#### EdU-seq analysis

Sequencing reads were adapter-trimmed using TrimGalore (v0.0.6, Babraham Bioinformatics) and aligned to the mm10 mouse and dm6 drosophila genome using bowtie2 (v2.5.3). Only uniquely mapping reads were kept using samtools (v1.15) and duplicates removed using picard MarkDuplicates (v2.27.3).

Coverage bigwigs and heatmaps were created with reference-adjusted reads per million (RRPM) spike-in normalisation using deeptools^135,136^. Relative genome-wide replication patterns were analysed by csaw^137^ using 5kb bins with normalisation by reads per million. Nearby adjacent significantly altered bins were clustered to find regions with different replication patterns. Nearby adjacent significantly altered bins were clustered to find regions where the distribution of replication was altered. Spike-in normalised log_2_ fold change of individual 5kb bins was obtained by normalising the raw read counts in each bin by per million Drosophila spike-in reads to obtain counts in reference-adjusted reads per million (RRPM) for each replicate. The mean RRPM count of three replicates from each bin was calculated for both dTAG and DMSO-treated samples, and these were used to calculate the log_2_ fold change of individual bins. Feature overlap was performed using GenomicRanges findOverlaps to extract bins overlapping with each feature. Initiation zones were obtained from^15^. Gene expression levels were obtained from control DMSO-treated TTchem-seq samples in this study. Repli-seq data was obtained from^138^ and replication timing was split into quintiles from earliest to latest replicating regions.

#### TTchem-seq analysis

TTchem-seq reads were trimmed with cutadapt (version 4.2) and mapped to a hybrid mouse (mm10) and Saccharomyces Cerevisiae (SacCer3) genome using STAR (version 2.7.11a) with special parameters’--alignIntronMax 1 --alignEndsType EndToEnd --winAnchorMultimapNmax 200 --outFilterMultimapNmax 100’ . Deduplication was performed with umitools (version 1.1.4). Fastq files were filtered for rRNA reads using ribodetector (version 0.3.1), and as a second round of filtering, reads mapping the the mouse rRNA sequence were also removed. Quantification of genes and repeats was performed with the TEcount tool, part of the TETranscripts suite (version 2.2.3). Differential expression analysis was performed with DeSEQ2^139^ using scale factors derived from the number of unique reads mapping to the SacCer3 genome. Expressed genes were defined as having a minimum of 10 unique reads in at least 3 replicates of each timepoint. Significant genes were defined as having a FDR < 0.01 and a minimum log_2_ fold change of 1. RPKM values for each gene were calculated using the mean read counts across the gene in DMSO conditions. Genes were split into quartiles based on RPKM levels in the DMSO samples to obtain annotations of Q4: high, Q3: mid-high, Q2: low-mid, and Q1: low transcribed genes. Genes without enough reads to be included for differential expression analysis were considered not expressed. RRPM values were calculated by expressing read counts per million spike-in reads and bigwig files were created using this normalisation with deeptools^135^. Spike-in normalisation of bigwig files was performed using the same scaling factors used for differential expression analysis. Transcription velocity was estimated by taking the RRPM normalised TTchem-seq signal in 50 bp bins and dividing by the spike-in normalised Pol II ChIP-seq signal in each bin.

#### RASAM

Sequencing reads were processed using PacBio software and custom scripts as described in^68^ to obtain the genomic localisation of single molecules and methylation probabilities and footprint widths and locations across each molecule. BrdU predictions were performed using custom scripts and models described in^67^ to obtain BrdU probabilities for each molecule. Molecules with a probability > 0.3 were classed as BrdU+. Nucleosome density for each fibre was calculated as in^140^ by estimating the number of nucleosomes present in each inaccessible region. The widths used are shown in Supplementary Figure 4D, and molecules containing inaccessible regions of size 11+ where nucleosome count could not be accurately estimated were filtered out. Autocorrelation analysis was performed as in^68^ for molecules over 1 kb in length. Autocorrelations were then clustered by Leiden clustering^141^ using scanpy with a resolution of 0.4. Clusters containing less than 10% of the total reads were removed, leaving 6 chromatin fibre types. Fisher’s exact tests were used to compare the enrichment of fibre types overlapping different genomic regions or in BrdU+ fibres. P-values were converted to q-values by FDR correction. Analysis over specific regions was performed by extracting the portion of the read overlapping any region and extracting nucleosome densities, footprint widths, and linker length over the region. Overlapping regions of less than 1 kb were excluded from downstream analysis. Linker lengths were calculated by measuring the length of the accessible region between two adjacent nucleosome-sized footprints (100-200bp). Gene expression quartiles were obtained from protein-coding genes from TTchem-seq data. H3K9me3 and H3K27me3 peaks were obtained from^25^. Fibre type enrichment over regions was obtained by extracting the fibre type cluster for reads where the first 1 kb used for autocorrelation analysis was within the target region. Single-molecule plots were obtained by selecting all reads which completely overlapped the desired window. Mean accessibility plots were created by calculating the mean accessibility at each base-pair across all molecules and smoothing this within a 33 bp rolling window. To assess chromatin repeat lengths, the portion of each read overlapping the desired regions was extracted and overlapping regions of less than 1 kb filtered out. The overlapping portion of each read representing a binary vector of accessible or inaccessible bases was used as input to a fourier transform using scipy.signal.periodogram to give a power spectral density estimation across a vector of frequencies. The inverse of the frequency vector then gives the repeat length in base pairs. Plots shown represent the summary of all spectral densities calculated across all overlapping reads

#### IDLI

Iteratively defined lengths of inaccessibility (IDLI) was performed on RASAM data as in^69^. Briefly, nucleosome footprints were called using increasing t probabilities (31, 51, 71) in the hidden Markov model to obtain sub-nucleosome footprints. The original footprints from RASAM data obtained using a transition probability (t) value of 1 were used as the nucleosome footprints. For nucleosome footprints <200bp in length, the accessibility data across a 140 bp window centred on the footprint midpoint was obtained for each t value (31, 51, 71). This was then used as the input for Leiden clustering with resolution 0.5. Clusters adding up to less than 5% of the total number of nucleosomes were filtered out. Footprint sizes were assessed by using the width of the nucleosome footprint (t=1) for each nucleosome in the cluster. Accessibility profiles relative to the footprint midpoint were calculated using t=71 values. Horizon plots were obtained by computing the z-scores of enrichment for subnucleosome footprint sizes and positions relative to the nucleosome midpoint. For visualisation, the z-scores were capped at 95% of the maximum value. Unmethylated footprints were filtered out before further analysis. Clustering was performed in R using the dist function with Euclidian distance and Ward.D2 clustering using hclust. Differential enrichment of nucleosome types was performed using Fisher’s exact test, and p-values corrected by FDR.

#### ChIP-seq analysis

ChIP-seq samples were sequenced paired-end and adapter-trimmed using CutAdapt (v4.2) and aligned to a hybrid mm10 mouse reference genome the dm6 drosophila genome using bowtie2 (v2.5.3). Samtools (v1.15.1) was used to filter unique reads and UMI deduplication was performed using umi-tools (v1.1.4). H2BK120ub1 ChIP-seq samples were processed without UMI deduplication and were deduplicated by Picard MarkDuplicates (v2.27.3). Reads were filtered against the ENCODE blacklist^142^ and a minimum mapping quality of 20 was used for downstream analysis. Replicate similarities were assessed by comparing the signal in 5kb bins genome-wide and replicates were averaged for making bigwig files. Spike-in normalisation factors were computed per million spike-in reads for each sample, correcting for differences in input spike-in percentages. Heatmaps and coverage profiles were made using deeptools (v3.5.5), using the average RRPM values from three replicates in each 50bp bin.

#### Micro-C analysis

Micro-C datasets were processed the distiller pipeline, with the original parameters for mapping, parsing, deduplication control, and binning as the yml in the repository (https://github.com/mirnylab/distiller-nf). Reads were mapped to the mouse reference assembly (mm10) using BWA-MEM (Burrows-Wheeler Alignment for 70 bp to 1 Mbp reads) with flags for high accuracy mapping of paired end reads (-SP). A chunk size of 30M reads was applied for optimized distribution computing.

Paired-end reads were parsed into read pairs using the pairtools package (’make_pairsam’, ‘drop_readid’, ‘drop_seq’), with additional options to store MAPQ scores (‘--add-columns mapq’) and mask complex walks in long reads (‘--walks-policy mask’). Reads considered PCR and optical duplicates, read pairs with a maximum allowed mismatch of 1 bp between pairs on both sides, were removed.

The processed read pairs were binned into multi-resolution contact maps (multi-cool/mcool files) using the cooler package, with resolutions ranging from 10 Mb to 100 bp (500 bp, 1 kb, 2 kb, 5 kb, 10 kb, 50 kb, 100 kb, 250 kb, 500 kb, 1 Mb). All contact matrices were normalized using the iterative correction procedure^143^, and pairs with low mapping quality, with MADmax (maximum allowed median absolute deviation) threshold of 30, were excluded during binning. The output files were balanced to ensure proper normalization of contact matrices.

The different biological and technical replicates were pooled into treatment groups (DMSO, dTag1h, dTag2h and dTag4h) using the library_groups section of the yml in distiller.

#### Cis/Trans read analysis

For the boxplot analysis of trans read counts in the non-titration data, we used mapped read files for each replicate generated by the distiller-nf pipeline. We first counted the reads for each nucleosomal orientation in cis and trans separately. To exclude non-ligated chromosomes, we discarded paired-end reads with forward-reverse (FR) nucleosomal orientation and summed the remaining orientations for cis, trans, and total reads without FR. We calculated the percentage of trans reads as (trans reads without FR / total reads without FR) * 100 and plotted the data points and boxplots for all replicates across treatments. We applied a Wilcoxon test to determine if the percentage distribution of trans reads for each treatment were significantly different from the DMSO control, resulting in a p-value of 0.014 for the DMSO vs. dTAG 4h comparison. Applying a t-test, which assumes a normal distribution, resulted in a p-value of 0.073 for the same comparison.

#### Eigenvector analysis

We computed the compartment profiles using the cooltools package^144^. First, we calculated the first principal component (PC1) for both the experimental samples and the control datasets from the multi-resolution cooler output from the distiller-nf pipeline. The eigenvector calculations were computed using the cooltools.eigs_cis function at multiple resolutions: 5 kb, 10 kb, 50 kb, 100 kb, and 500 kb. Phasing of the eigenvectors was done based on GC content to ensure accurate A/B compartment identification. To visualize the distribution of the read densities for each sample, we used a custom R script to generate density plots with ggplot2 library.

### Microcompartment detection

To identify microcompartments, we used the eigenvector one derived from A/B compartment analysis, but with higher resolution data 5 kb resolution instead of the 50 kb resolution. We selected the 5 kb resolution as it provides a clear visualization of the microcompartments in HiGlass, where they are represented by more than one consecutive pixel. By computing the eigenvector one with cooltools^144^ through principal component analysis (PCA), we captured the most variance in our data. This eigenvector represents the greatest changes and direction of change from our data at higher resolution, analogous to how A/B compartments are quantified at 50 kb resolution.

To identify microcompartment loci, we calculated the differential between DMSO and the most extreme treatment, FACT dTAG 4h, where microcompartments were more pronounced. We plotted the differential for each chromosome, with the x-axis representing the chromosome coordinate and the y-axis representing the differential value (dTAG4h-DMSO). Visual inspection of the differential revealed that the values were consistently spread along the x-axis, indicating good coverage across both samples along the chromosome. However, for some chromosomes, the differential values started at higher levels near the beginning of the chromosome and decreased as the distance progressed, or vice versa, forming a gradient-like pattern.

Since PCA is calculated independently for each chromosome, we applied independent corrections to these distributions. Specifically, we centred eigenvalue differential to 0 in bin windows of 5% of the total chromosome length. This method preserved the vertical distribution of the original differential, including the extreme values, while providing a general threshold across the entire chromosome. We then saved the loci of the 5 kb bins where the differential values were at the most extreme positive and negative thresholds. These thresholds were applied independently for each chromosome at 5%, 2.5%, 1.25%, and 0.625% most positive and most negative levels.

We visually inspected the overlap of these regions with microcompartments in HiGlass. No obvious features overlapped with the negative thresholds, while positive thresholds showed consistent overlap with microcompartments. The top 0.625% positive values included more false negatives, and the 5% threshold had more false positives. Therefore, the 2.5% threshold was deemed the most suitable, as it showed the best overlap with microcompartments when visually inspected on chromosomes 1, 2, 7, and 11, as well as other regions previously associated with microcompartments. However, we also used the other thresholds for further downstream analysis to examine the gradient of stringency. For each threshold, we generated BED files for the microcompartment regions using two different approaches. In the first, we retained regions strictly from the 5 kb bins, while in the second, we merged contiguous regions with a 2-bin spacing (10 kb) between them.

### Loops and compartment pile-ups

Loop and compartment pile-ups we generated with cooltools and coolpuppy according to “walk-through” notebooks (https://cooltools.readthedocs.io/en/latest/notebooks/pileup_CTCF.html) and (https://coolpuppy.readthedocs.io/en/latest/Examples/Walkthrough_CLI.html#rescaling). For CTCF intersections we computed from annotated CTCF peaks (GSM2418860). For enhancers, FANTOM5 annotation of active enhancers in the mm10 genome (https://dbarchive.biosciencedbc.jp/data/fantom5/datafiles/reprocessed/mm10_latest/extra/en hancer/). For promoters, the AMC annotated gene list was used. The read occupancy at each site was computed for each list from DMSO Micro-C data (read1). The top and bottom 1% of sites were excluded from analysis to reduce outliers.

### Machine Learning-Based Microcompartment Feature Analysis

ChIP-seq occupancies were computed in dTAG-depleted samples using csaw^136^ RegionCounts across each expressed gene obtained from TTchem-seq data. SUZ12, RING1B, and RAD21 occupancies were computed within a 1 kb window centred on the TSS. H3K27ac, H3K4me3, and H2A.Z occupancies were computed within a 2.5 kb window centred on the TSS. H2BK120Ub1, H3K36me3, H3.3, and RNA Pol II occupancies were computed across the gene body. H2B and H3 occupancies were computed in a window between the TSS and 2.5 kb downstream to assess the effects of TSS proximal histone loss. ChIP counts at each gene were filtered for enrichment over the input by a minimum of log_2_(1.58) except for histone H2B and H3 ChIPs, which used an enrichment threshold of log_2_(0). Occupancies were computed in RRPM^135^ and values used were the mean of n=3 biological replicates. Genes without enrichment over input for a specific ChIP were set to a value of 0. Gene body and promoter GC content were calculated with gcContentCalc from Repitools using the whole gene body or a window of -3000 bp to +100 bp centred on the TSS respectively. The number of genes within 1 Mb represent the number of expressed protein-coding genes within 1 Mb of each gene. Exon density was calculated as exon counts per kb. Distance to IZ was calculated using the minimum distance to an initiation zone obtained from^58^.

For AMC prediction modelling, all features were extracted and combined with the AMC threshold of the gene. AMC-overlapping genes representing the 5% to 2.5% threshold were excluded from the training data due to the high false positive rate within these genes. H3 and H2B occupancy and SUZ12 and RING1B occupancy were found to be highly correlated (pearson correlation > 0.8) and were therefore merged into nucleosome occupancy (H2B and H3) and polycomb repressive complex (PRC, SUZ12 and RING1B) respectively. 16 cross-validation splits were created using 4 chromosomes as the test set and the rest as the training set, with all chromosomes being sampled in the test set at least once. Chromosome X and Y were excluded from the analysis. AMC overlapping genes were assigned a dummy variable of 1 as the prediction label. A boosted random forest model was trained using xgboost with the parameters: max.depth = 4; eta = 0.1; nrounds = 70; gamma = 0.01; objective = ‘binary:logistic’; booster = ‘gbtree’; num_parallel_tree = 1. Weights were calculated to account for any class imbalances in the training set by using a ratio of the sum of negative instances to the sum of positive instances and this was used in the scale_pos_weight parameter. Predictions were classified as AMC if the prediction probability was >0.5. Performance metrics were calculated using caret confusionMatrix and SHAP values were calculated using SHAPforxgboost. Clustering of SHAP values was performed using the 10 most important features by hierarchical clustering with the ward.D2 method and Euclidean distance. The number of clusters was determined by the silhouette method.

### Expression-Matching Gene Sets

To control for di9erences in gene expression when analyzing aberrant microcompartments (AMCs), we performed expression-matching using the MatchIt package in R. Genes were classified as containing AMC or not based on the top 0.625% most positive values on the di9erential eigenvector values (see Microcompartment detection section in Methods). The original dataset included 1761 AMC genes and 14104 non-AMC genes. We then matched AMC genes to non-AMC genes with similar expression levels in the DMSO control condition. Matching was carried out using nearest neighbor matching with a 1:1 ratio, without replacement, and a caliper of 0.1 to restrict matches to genes with comparable expression. This procedure yielded two balanced sets of 1723 AMC and 1723 non-AMC genes, allowing us to isolate the e9ects associated with AMC status independently of baseline expression levels.

### Antibodies

**Supplementary Table 1.**
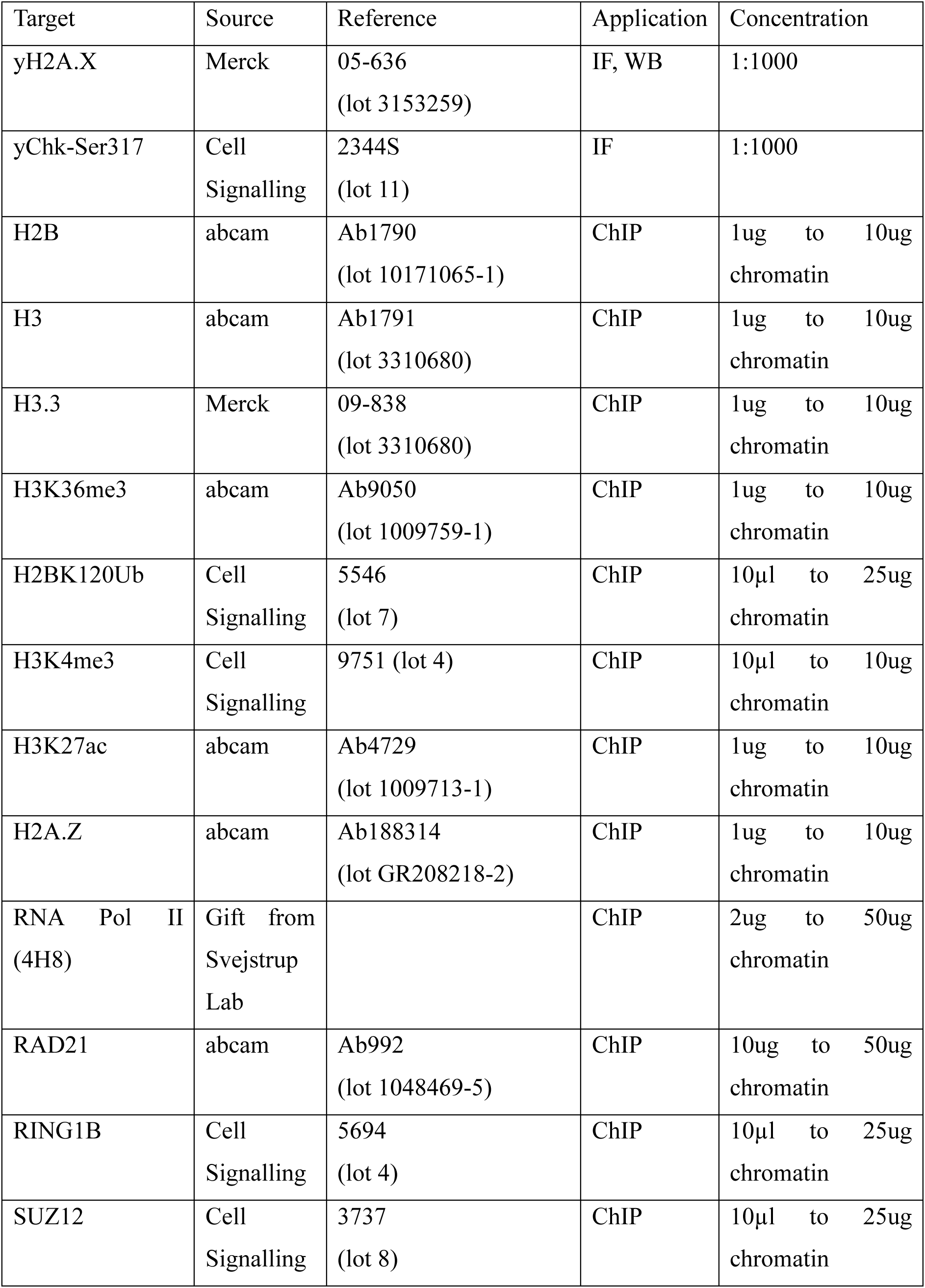
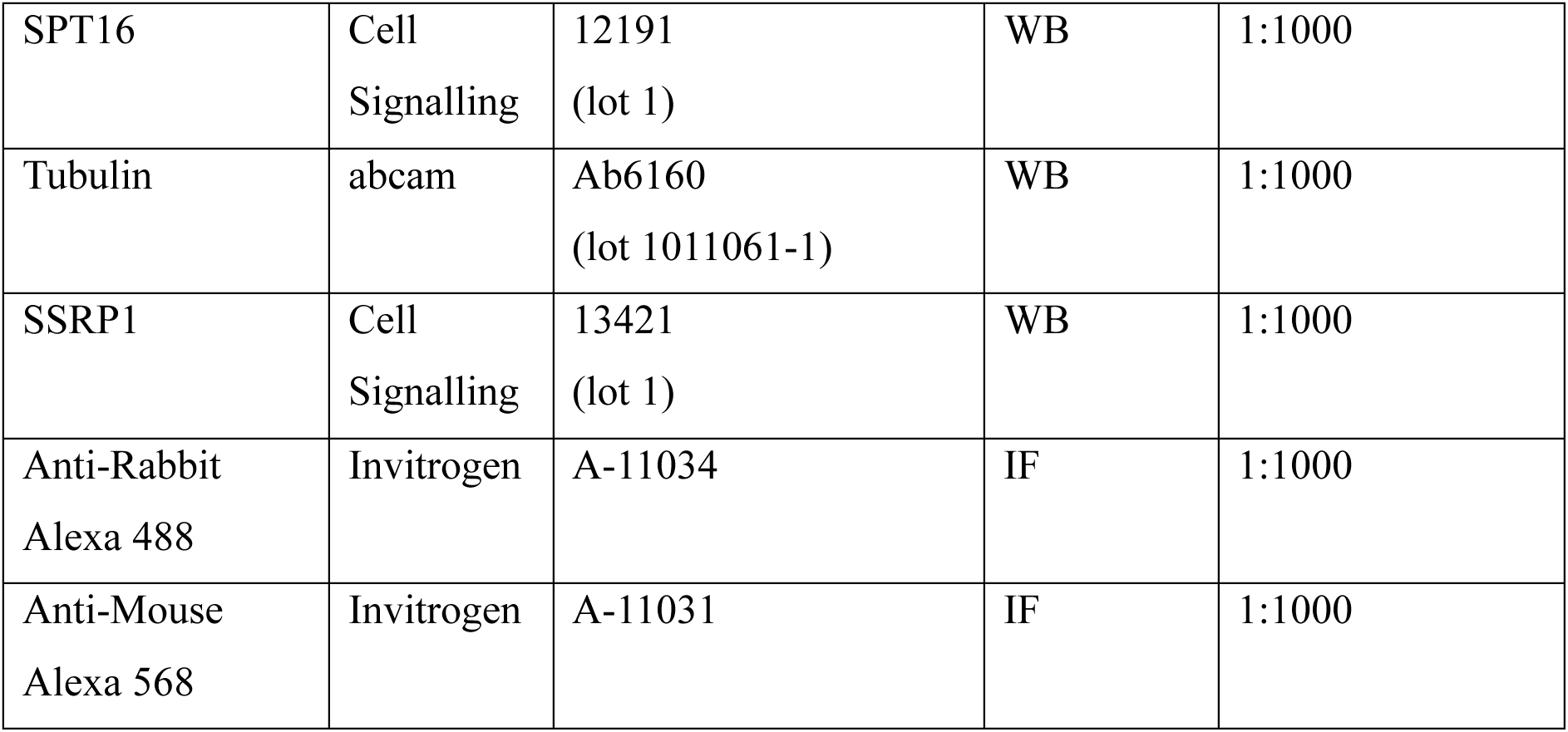
Antibodies used in this study and applications used for (WB: western blot, IF: immunofluorescence, ChIP: chromatin immunoprecipitation)

**Supplementary Table 2.**
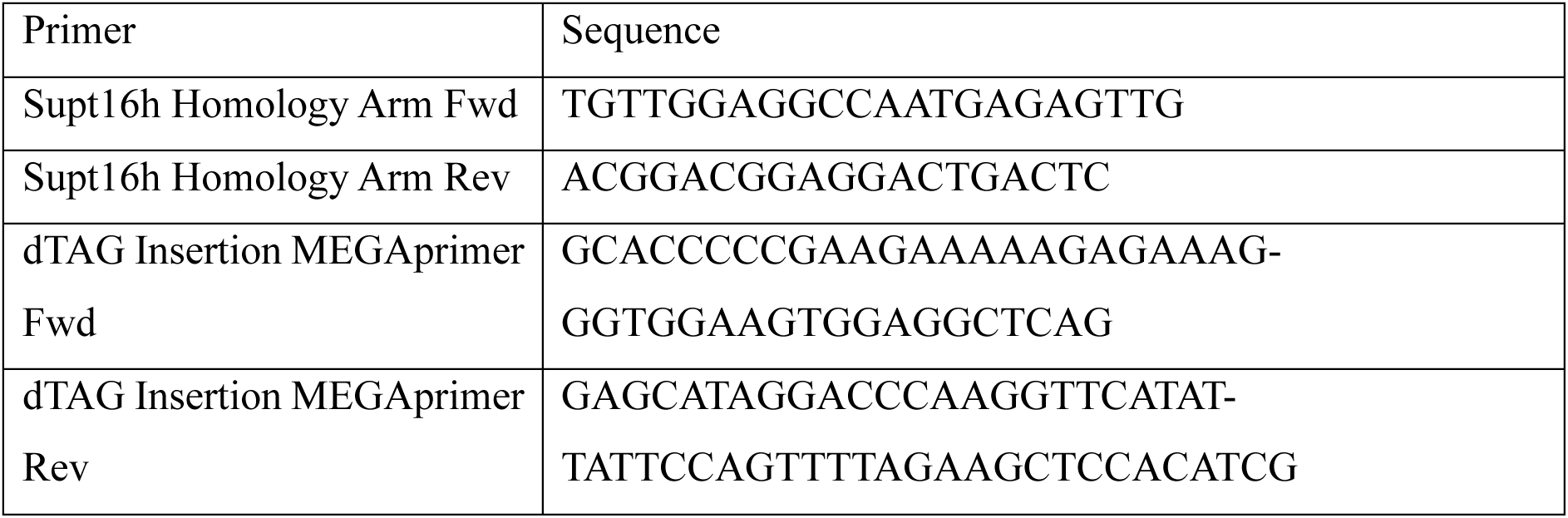
Primer sequences used in this study.

## Supplementary Figures

**Figure S1.**
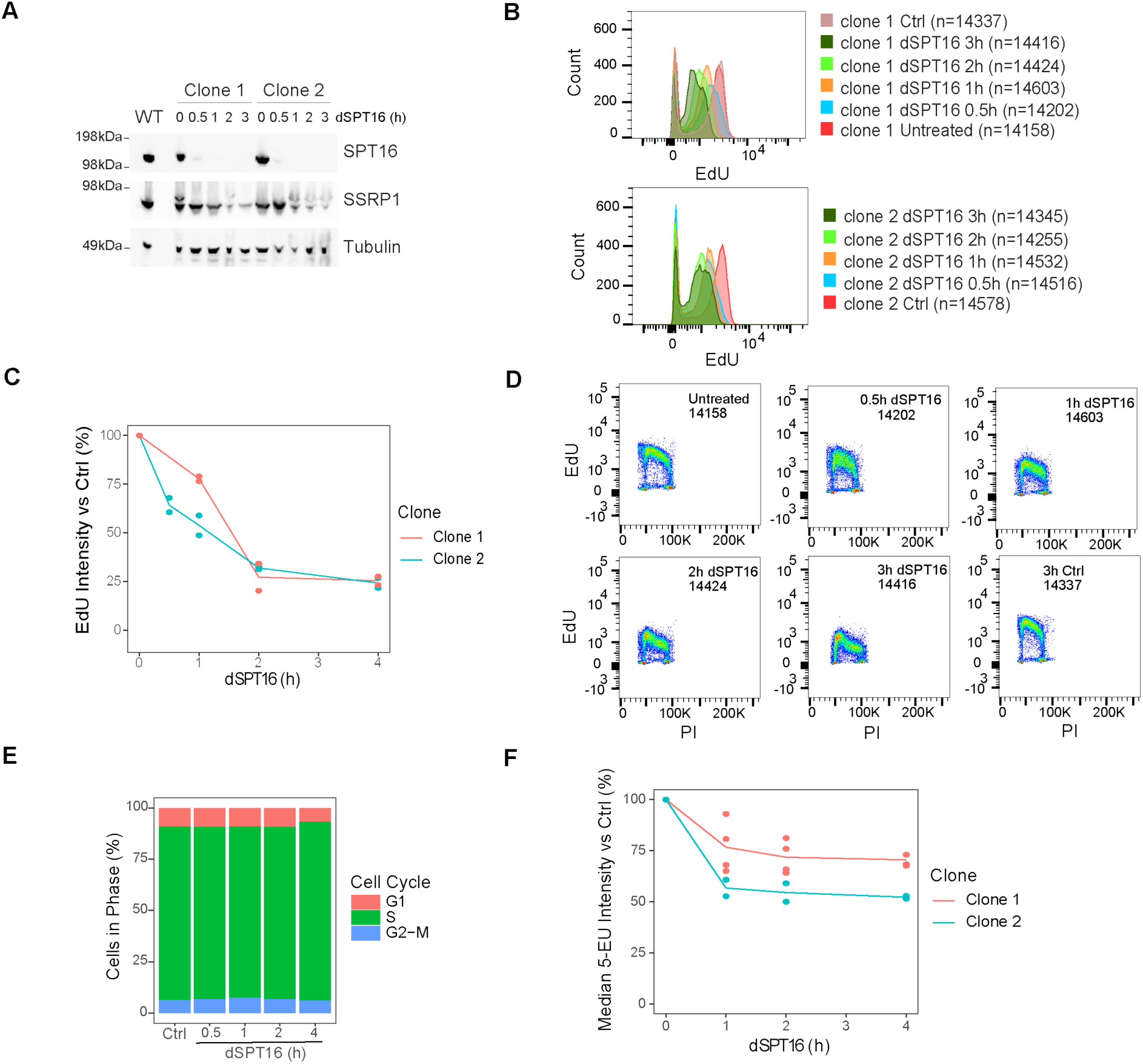
SPT16-dTAG mESC Validation Related to Figure 1. **A.** Western blot of SPT16, SSRP1, and α-Tubulin in two monoclonal SPT16-dTAG mESC cell lines after treatment with 1 μM dTAG-13 (dSPT16) for the specified time. Parental E14 Ju mESCs (WT) are also shown. **B.** 5-ethynyl deoxyuridine (EdU) incorporation levels in two clonal cell lines with SPT16-dTAG treated by dSPT16 or DMSO control (Ctrl) assessed by flow cytometry **C.** EdU incorporation changes in two clonal SPT16-dTAG cell lines after dSPT16 for the specified time in hours (h). Values are shown relative to Ctrl cells. Each point represents a distinct biological replicate (n=2 per clone) **D.** Flow cytometry profiles of EdU incorporation and total DNA content by propidium iodide (PI) showing the cell cycle of SPT16-dTAG mESCs (Clone 1) treated for the specified times with dSPT16. DMSO vehicle (Ctrl) and untreated controls are also shown. **E.** Proportion of cells from microscopy in Figure 1C in each cell cycle phase in Ctrl and dSPT16 cells (Clone 2). **F.** Median 5-ethyl uridine (5-EU) incorporation relative to Ctrl cells of SPT16-dTAG mESCs treated with dSPT16 for the specified time. Biological replicates in two clonal cell lines are shown (n=4 Clone 1, n=2 Clone 2).

**Figure S2.**
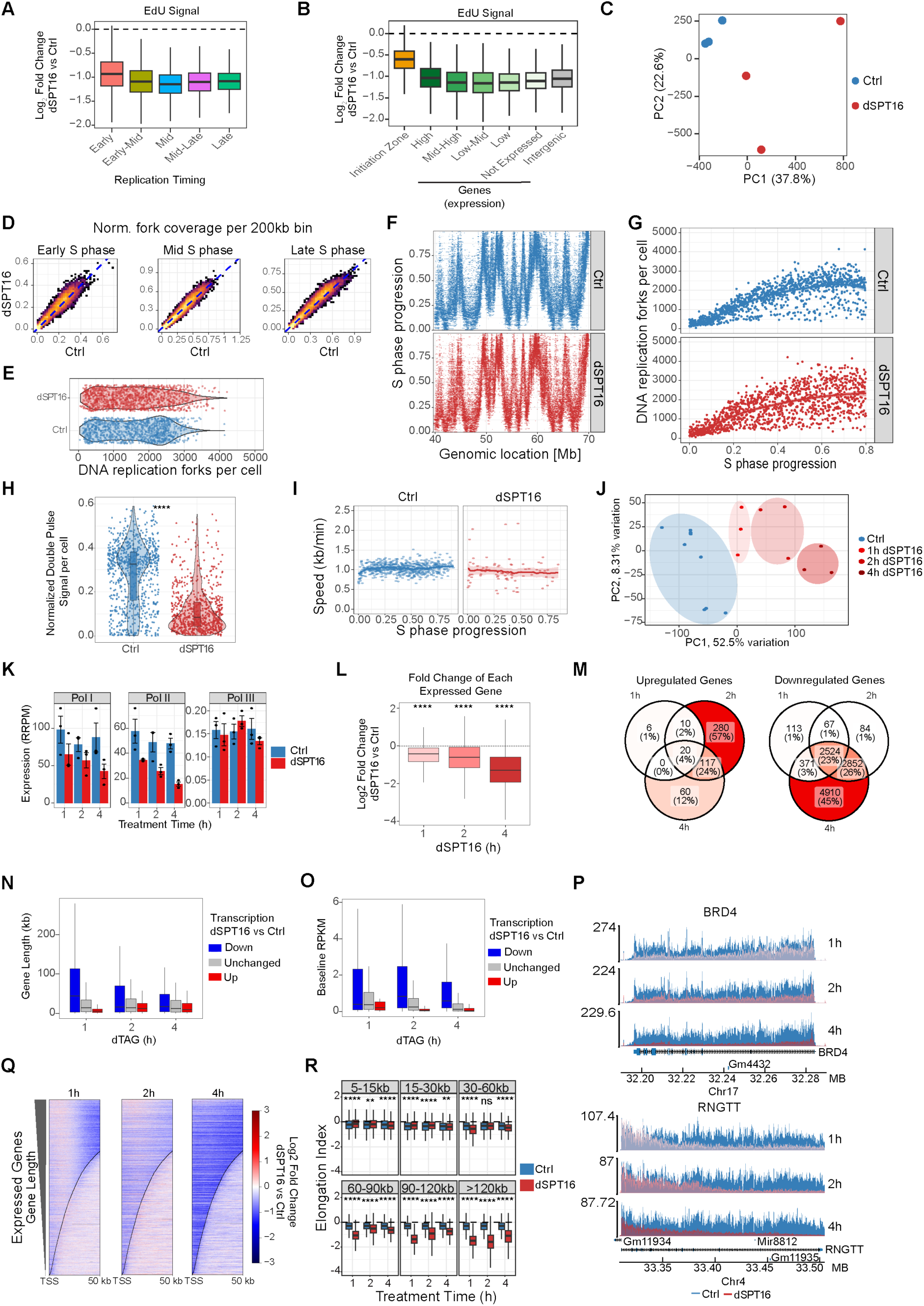
EdU-seq Validation and TTchem-seq Transcriptional Changes Related to Figure 2. **A.** EdU signal quantified by spike-in (reference-adjusted reads per million (RRPM)) in SPT16-dTAG mESCs. Shown is log_2_ fold change of EdU signal in 1 hour dTAG-treated cells (dSPT16) versus DMSO control (Ctrl) analysed in 5kb bins genome-wide. Regions were stratified by replication timing quintiles from earliest to latest replicating across S phase. **B.** EdU signal (as in B.) in 5kb bins overlapping IZs, genes grouped by expression quintile from TTchem-seq, or intergenic regions. **C.** PCA of genome-wide replication signal from EdU-seq. **D.** Replication timing correlation between 1h dSPT16 and Ctrl for the replication patterns of cells in early, mid, or late S phase detected by DAPI staining. **E.** Number of replication forks detected per cell by scEdU-seq in dSPT16 and Ctrl cells. **F.** Genomic localisation of detected replication forks in single cells across a single chromosome. Y-axis shows the replication timing status determined by DAPI intensity. **G.** The number of detected replication forks for single cells across S phase progression assessed by DAPI. **H.** Double pulse signal (measuring the ratio of the strength of the second EdU signal relative to the first) for single cells. **I**. Replication speed in single cells for double-pulse positive cells in which replication fork speed could be calculated. **J**. PCA of global gene expression assessed by nascent RNA sequencing (TTchem-seq) in SPT16-dTAG mESCs treated for 1, 2, or 4 hours with dSPT16 or 1, 2, or 4 hours DMSO control (Ctrl) in n=3 biological replicates per condition. **K.** Spike-in normalised (RRPM) read coverage at Pol I, Pol II, and Pol III genes in dSPT16 or control cells treated for the specified times. **L.** Log_2_ fold change of all expressed genes after dSPT16 treatment for the specified time. Significance was calculated by one-sample t-test against zero (****: p < 2.2x10^16^). **M**. Number of differentially expressed up- or down-regulated genes shared after different lengths of dSPT16 treatment. **N.** Length of differentially expressed (up or down) or not significantly altered (Unchanged) genes after dSPT16 for the specified time. **O.** Baseline expression levels (calculated as reads per kilobase-million (RPKM) in Ctrl samples) of differentially expressed (up or down) or not significantly altered (Unchanged) genes after dSPT16 as in N. **P**. Single gene tracks of spike-in quantified (RRPM) TTchem-seq signal over two expressed long genes. Gencode annotation shown below. **Q**. Heatmap of change in TTchem-seq signal after dSPT16 across all expressed genes from the transcription start site (TSS) to 50kb downstream. Genes are sorted by descending length and the dotted line represents the transcription end site of each gene. **R**. Elongation index of all genes after dSPT16 or Ctrl treatment. Elongation index represents the ratio of TTchem-seq signal in the first 5% of the gene body vs the last 5%. Significance was calculated using paired t-test (NS: not significant, * p<0.05, **p<0.01, ***p<0.001, ****p<0.0001).

**Figure S3.**
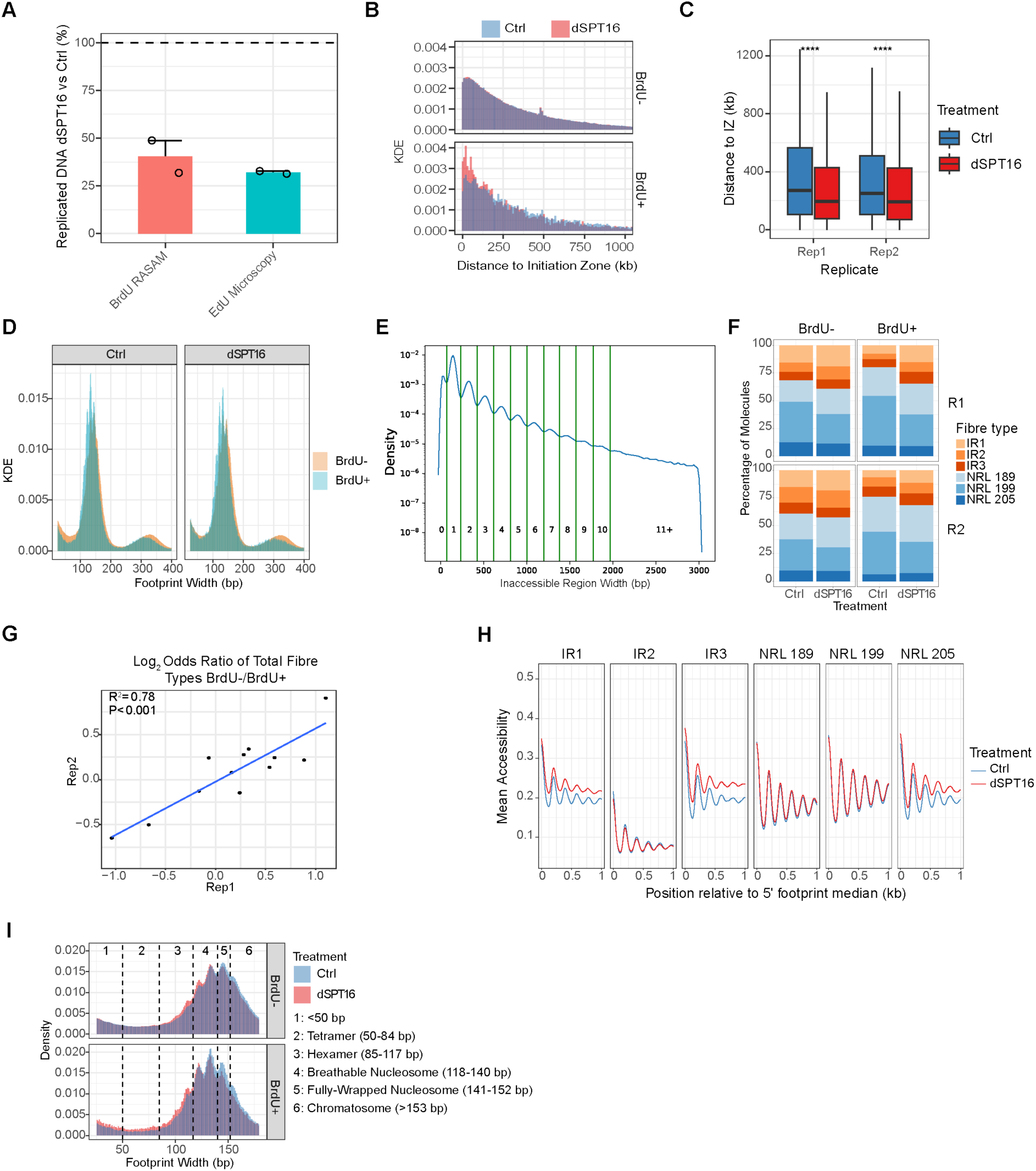
RASAM Validation Related to Figure 3. **A.** Relative number of BrdU+ fibres detected by RASAM or EdU signal assessed by microscopy in 3 hour dTAG-treated (dSPT16) SPT16-dTAG mESCs compared to DMSO control (Ctrl). **B,C.** Distance to the nearest initiation zone (IZ) in chromatin fibres after dSPT16 or Ctrl treatment. Differences were calculated by Wilcoxon ranked-sum test (**** p < 0.001). **D.** Distribution of footprint widths in nascent (BrdU+) and bulk (BrdU-) fibres from cells treated with dSPT16 or Ctrl. **E.** Schematic for the estimation of nucleosome count on single fibres. Cutoffs were defined to estimate the number of nucleosomes within an inaccessible region based on the distribution of inaccessible region lengths. Green lines represent the cutoffs and the numbers represent the number of estimated nucleosomes within each inaccessible region. **F.** Proportion of chromatin fibre types in bulk (BrdU-) and nascent (BrdU+) fibres after dSPT16 or Ctrl treatment in two biological replicates (R1 and R2). **G.** Correlation of log_2_ odds ratios for enrichment of fibre types in dSPT16 fibres between two biological replicates. Pearson correlation and p value are shown. **H.** Mean accessibility across the first 1 kb of different fibre types after dSPT16 or Ctrl treatment. **I.** Footprint widths in dSPT16 and Ctrl bulk and nascent fibres. Size ranges of different nucleosome classes are shown.

**Figure S4.**
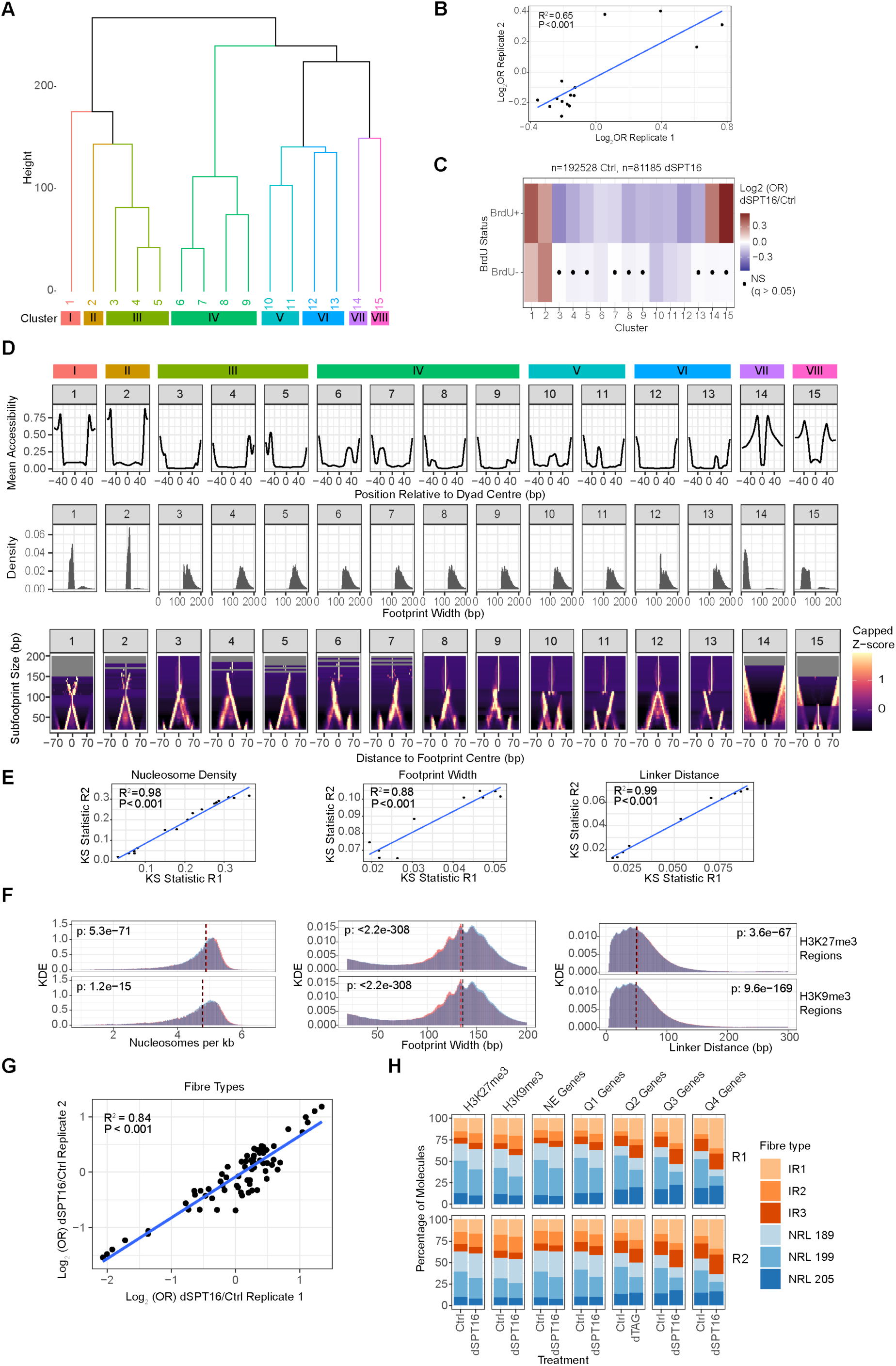
IDLI clustering and SAMOSA validation Related to Figure 3. **A.** Dendrogram of distances between each nucleosome type for clustering. **B.** Correlation of log_2_ odds ratios for enrichment of nucleosome types on nascent (BrdU+) fibres in dTAG-treated SPT16-dTAG mESCs (dSPT16) compared to DMSO-treated control (Ctrl). **C.** Heatmap of enrichment of nucleosome types in bulk (BrdU-) and nascent (BrdU+) fibres after dSPT16. Significance was calculated by Fisher’s exact test, and dots indicate nucleosome types that were not significantly altered. **D.** (top) Mean accessibility across each nucleosome type. (middle) distribution of footprint widths within each nucleosome type. (bottom) V-plots of subfootprint sizes and distance relative to the footprint centre, showing accessibility patterns across each nucleosome type. Z-score shown is capped at the 95% percentile of values. **E:** Correlation of Kolmogorov-Smirnov statistics (measuring the difference between dSPT16 and Ctrl) between two biological replicates for differences in nucleosome density, footprint width, and inter-nucleosome linker distance. Pearson correlation coefficient and p-value are shown. **F.** Distribution of nucleosome density, footprint width, and linker distance in dSPT16 or Ctrl fibres overlapping H3K27me3 or H3K9me3 regions. P-values from Kolmogorov-Smirnov test are shown. **G.** Correlation between two biological replicates of log_2_ odds ratios of fibre type enrichment in dSPT16 fibres across annotations from Figure 3I. Pearson correlation coefficient and p-value are shown. **H.** Proportion of each fibre type for dSPT16 and Ctrl chromatin fibres overlapping each annotation in two biological replicates (R1 and R2).

**Figure S5.**
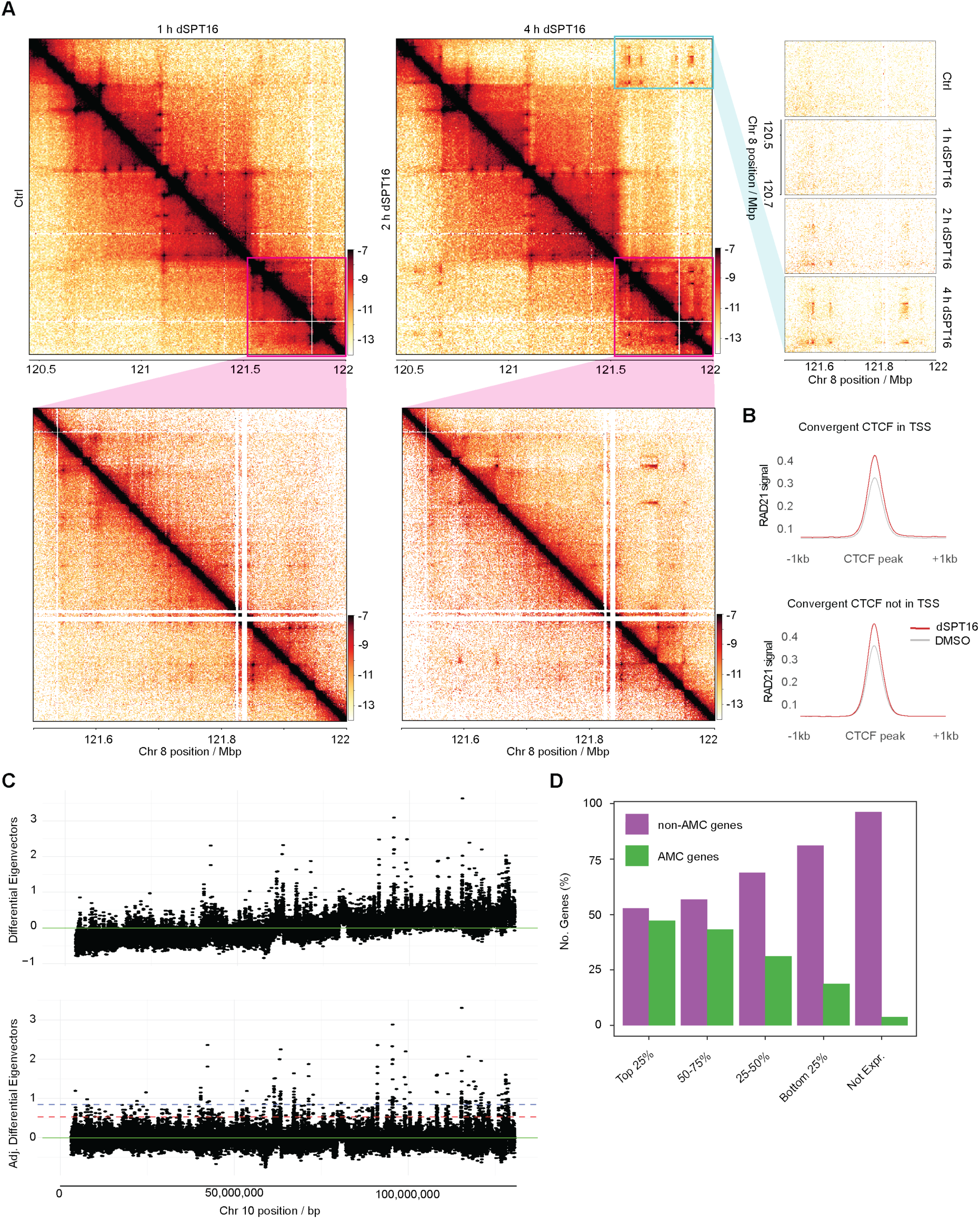
Related to Figure 5. **A.** Micro-C interaction heatmaps of gradually increasing AMCs between distal AMC forming regions upon SPT16 depletion by dTAG (dSPT16) compared to DMSO control (Ctrl). Top, left panel: Ctrl (bottom, left), 1 h dSPT16 (top, left), 2 h dSPT16 (bottom, right), and 4 h dSPT16 (top, right) at 10 kbp resolution. Bottom, left panel: same as top panel, but zoomed into one AMC forming region at 5 kbp resolution. Right panel: Off-diagonal zoom between distal AMC regions of Ctrl, 1, 2, and 4 h dSPT16 conditions (top to bottom). **B.** RAD21 qChIP signal at CTCF sites stratified by proximity to TSS (top) or not (bottom). **C.** Differential Eigenvalue before (top) and after (bottom) local adjustment. Green line: 0 position. Red line: 2.5% threshold. Blue line: 0.625% threshold for AMC calling. **D.** Fraction of genes stratified falling into AMCs by expression quartiles determined by TTchem-seq in Ctrl cells.

**Figure S6.**
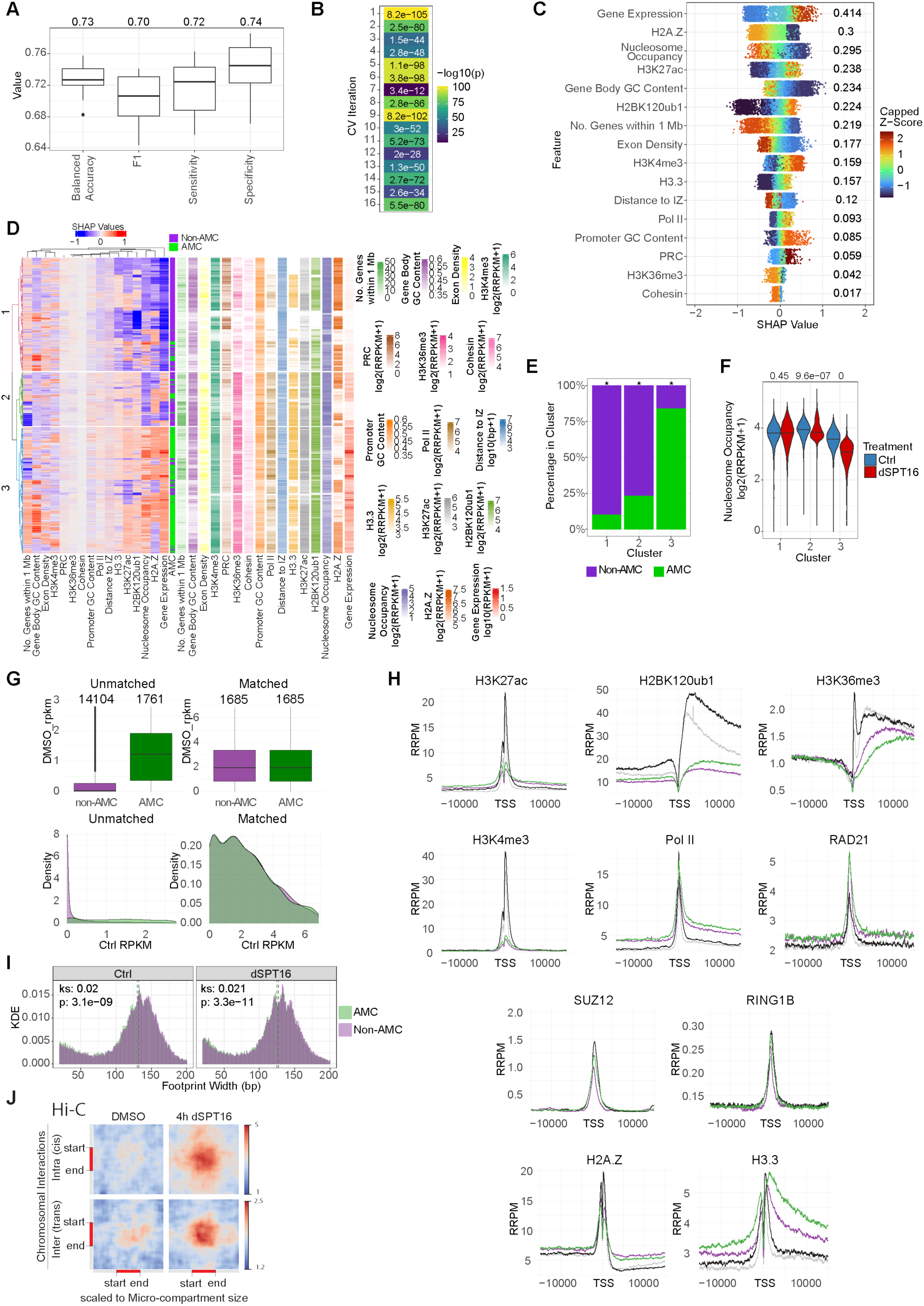
Related to Figure 6. **A.** Balanced accuracy, F1 score, sensitivity, and specificity of the AMC prediction models across n=16 cross-validation folds. Mean values across all cross-validation folds are shown. **B.** Heatmap of p-values of prediction model accuracy against a random classifier for each cross-validation iteration. **C.** Mean SHAP values across cross-validation iterations for all correctly-classified genes (n=7365). Each gene is coloured by the feature value normalised by z-score. Z-scores for each feature were capped such that the maximum and minimum values represent the 90^th^ and 10^th^ percentiles respectively. **D.** Hierarchical clustering of correctly-predicted genes based on SHAP values. AMC overlap and feature values for all features included in the model are shown. **E.** Bar plot showing the percentage of AMC overlapping and non-overlapping genes in each cluster. Enrichment was calculated by Fisher’s exact test (*: p < 2.2x10^16^). **F.** Spike-in normalised (reference-adjusted reads per million: RRPM) H3 occupancy in the first 2.5 kb of the gene body for genes in each cluster after 4 hours of DMSO (Ctrl) or dTAG (dSPT16) treatment. Differences were tested by Wilcoxon ranked-sum test and p-values are shown. **G.** Gene expression (expressed as reference-adjusted reads per kilobase million (RPKM) in Ctrl cells) of AMC-overlapping and non-overlapping genes before and after expression matching. **H.** Spike-in normalised ChIP-seq occupancy of various targets at expression-matched AMC and non-AMC genes expressed as RRPM, reference-adjusted reads per million **I.** Footprint widths in AMC overlapping and AMC non-overlapping regions of partially overlapping AMC genes. Dotted lines represent the median values and Kolmogorov-Smirnov effect size (ks) and p-value are shown. **J.** Cis and trans chromosomal domain interaction pile-ups for AMCs from Hi-C.

## Notes

### Competing Interest Statement

The authors have declared no competing interest.

